# A threshold level of JNK activates damage-responsive enhancers via JAK/STAT to promote tissue regeneration

**DOI:** 10.1101/2024.10.31.621241

**Authors:** John W. Quinn, Mariah C. Lee, Chloe Van Hazel, Melissa A. Wilson, Robin E. Harris

## Abstract

Tissue regeneration requires precise activation and coordination of genes, many of which are reused from development. While key factors have been identified, how their expression is initiated and spatially regulated after injury remains unclear. The stress-activated MAP kinase JNK is a conserved driver of regeneration and promotes expression of genes involved in proliferation, growth, and cell fate changes in *Drosophila*. However, how JNK selectively activates its targets in damaged tissue is not well understood. We previously identified Damage-Responsive, Maturity-Silenced (DRMS) enhancers as JNK-activated elements critical for regeneration. Here, we show that cell death is dispensable for the activation of these enhancers, which only depends on JNK and its immediate downstream effectors. One of these is JAK/STAT, which acts as a direct, additional input necessary to expand enhancer activity into the wound periphery where JNK alone is insufficient. Furthermore, we demonstrate that a threshold level of JNK is required to initiate enhancer activation. Together, our findings reveal how JNK and JAK/STAT signaling cooperate to drive spatially and temporally regulated gene expression through damage-responsive enhancers, ensuring proper regenerative outcomes.

## Introduction

Regeneration, the repair and regrowth of damaged tissues, occurs across numerous species in diverse tissue contexts, though often poorly in humans (1–3). It has been well established that regenerative growth often reuses developmental signaling; for example, FGF8, retinoic acid and Sonic Hedgehog are crucial for both salamander limb development and for its regrowth following amputation, with striking similarities in the temporal and spatial expression patterns of each factor (4). Thus, regenerative failure likely stems not from an absence of necessary genetic factors, but the inability to coordinately reinitiate their expression after injury. Until recently it was unclear how a single gene could be expressed differently in the contexts of regeneration and development, but the discovery of damage-responsive enhancers have provided an explanation (5–7). These enhancers have been identified in diverse organisms and tissues, and often share traits like modularity, common signaling inputs and epigenetic regulation (5–7). Their study has already revealed key insights into the identity, initiation and behavior of regeneration-promoting genes, and enabled strategies to improve regeneration even in non-regenerative organisms (8–10). Advancing our understanding of these elements is therefore essential.

To study how regenerative gene expression is initiated and regulated, we use the *Drosophila* wing imaginal disc, a larval epithelial tissue that forms adult wing structures, as a model (11). Extensively used to investigate fundamental aspects of development (11, 12), the disc has also become a powerful platform for dissecting the genetics underlying regeneration (13–15). Our prior work using the wing disc identified discrete genomic regions that change in accessibility after damage, correlating with altered expression of nearby genes, suggesting they likely function as regeneration-associated enhancers (8, 16). Using a genetic ablation system that we developed called DUAL Control (DC) (16, 17), we focused on two regeneration-promoting genes, *wingless* (*wg,* the *Drosophila Wnt1*) and *Matrix metalloproteinase 1* (*Mmp1*). Both are strongly induced by damage in regeneration-competent early third instar (L3) discs, but only weakly in late L3 discs that fail to regenerate (8, 16). We identified enhancers at both loci with similar modular structures: a damage-responsive (DR) region activated by injury, and a maturity-silencing (MS) region that limits enhancer activity in older discs via epigenetic silencing. Together these modules form DRMS enhancers that drive damage-induced gene expression, which subsequently becomes limited in late L3 discs, even in the presence of activating signals (8). Importantly, their modularity allows independent study of the DR and MS regions, a fact we have taken advantage of here. DRMS enhancers are activated by JNK signaling directly via AP-1 sites (8). Using their altered epigenetic signature, we subsequently identified dozens of additional putative DRMS enhancers with similar features, suggesting coordinated regulation of multiple genes by this mechanism (16). JNK activation and epigenetic regulation appear to be core aspects of regeneration in wing discs (18), and for tissue repair via damage-responsive enhancers in other species (5, 6). Thus, understanding these enhancers has broader implications for regenerative biology.

Crucially, JNK signaling during normal development, such as in the embryo or wing disc notum, fails to activate these enhancers or their regenerative targets (8). This suggests that either an injury-specific environment allows JNK to induce enhancer activity, or that additional pathways are required, or both. To explore this, we have investigated the involvement of additional regulatory factors. We find that, while JNK is necessary for DR enhancer activation, cell death and its resulting downstream signaling events are not. Moreover, JAK/STAT, a target of JNK, is required for full enhancer activation and proper spatial expression. Finally, by isolating JNK from the feedforward mechanisms that arise following cell death, we show that distinct JNK thresholds control target gene activation: high JNK levels at the center of a wound are sufficient to activate DR enhancers alone, while lower levels in the wound periphery require the additional input of JAK/STAT. At the lowest levels of JNK, where JAK/STAT is not induced, only a limited subset of targets respond, such as those expressed during development like *puckered (puc)*,. Thus, DR enhancers integrate JNK intensities generated by a wound alongside JAK/STAT input to direct gene expression with spatial and temporal precision during regeneration.

## Results

### Cell death is dispensable for activation of the regeneration program via DR enhancers

To investigate how DR enhancers are activated during regeneration, we examined an enhancer at the WNT locus known to drive *wg* and *wnt6* expression upon injury (DRMS^WNT^, (8, 16, 19) (Fig 1A). We used a GFP reporter driven by the DR module of the enhancer (*DR^WNT^-GFP*) (Fig 1A), which is inactive in undamaged tissue and unaffected by developmental stage when isolated from the MS region (8) (Fig 1B). We induced damage using the DC ablation system (16, 17), which triggers apoptosis in the distal pouch via heat-shock induced expression of activated *hemipterous* (DC^hepCA^) (Fig 1C). Cell death activates the regeneration program, observable at ∼6 h after heat shock (AHS) until ∼48 h AHS, with significant regenerative processes represented at the 18 h time point (17) (Fig 1D). Unlike other ablation systems that express a cell death stimulus continuously over an extended 20 - 40 h period (20–22) the DC system generates an acute injury that separates ablation from regenerative processes, making it ideal for dissecting DR enhancer activation. The DC system also includes a heat-shock activated pouch GAL4 driver (*hs-FLP; DVE>>GAL4*) to manipulate gene expression in the surrounding regenerating cells (17)(Fig 1C-D).

**Figure 1:**
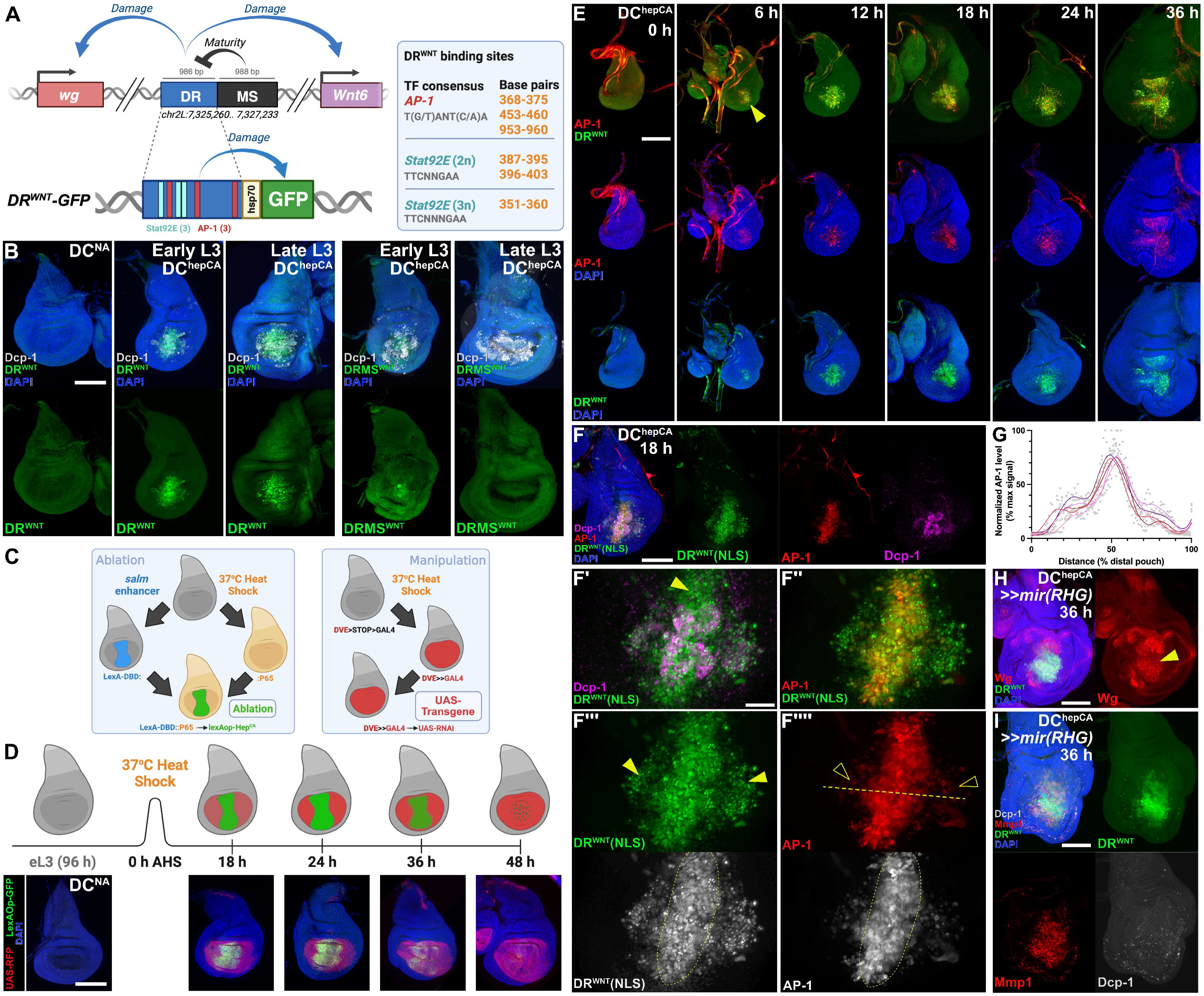
DR^WNT^ and the regeneration program can be activated independently of cell death. (**A**) Schematic of the DRMS^WNT^ regulatory element (top) with separable damage-responsive (DR) and maturity-silencing (MS) regions controlling *wg* and *Wnt6* expression. AP-1 and Stat92E binding sites in the DR region are noted (table, right). The DR^WNT^ reporter (bottom) comprises the enhancer driving GFP via an *hsp70* promoter. (**B)** DR^WNT^ and DRMS^WNT^ reporters in discs ablated with DC^hepCA^ at early (84h AED) or late (108h AED) L3. DR^WNT^ is activated in both stages, while DRMS^WNT^ is only active in early L3. Activation is not seen in non-ablating controls (DC^NA^). (**C**) The DUAL Control (DC) ablation system uses heat shock to activate LexA-based ablation in the distal pouch (green) and GAL4-driven gene manipulation across the pouch (red) (17). (**D**) Schematic of ablation timing with corresponding discs. *LexAop-GFP* (green) marks ablation domain; *UAS-RFP* (red) marks GAL4 activity. Ablation is transient, while GAL4 expression persists through regeneration. A non-ablating version of DC (DC^NA^) is used to show reporter dynamics. (**E**) Time course of *DR^WNT^-GFP* (green) and *AP-1-RFP* (red) reporters in DC^hepCA^-ablated discs shows overlapping activation from 6 to 36h AHS. (**F**) Disc with DR^WNT^[NLS]-GFP (green), AP-1-GFP (red), and Dcp-1 (magenta) at 18h AHS. (F’-F’’’’) DR^WNT^ activity occurs in cells lacking Dcp-1 (F’, yellow arrowhead). DR^WNT^ (F’’’, green) is active at the wound periphery, where AP-1 (F’’’’) is reduced. Dotted circle in (F’’’) and (F’’’’) indicates the observed separation of high and low JNK activity. Line in (F’’’’) shows plane of quantification in (G). (**G**) Quantification of AP-1 fluorescence profiles across wounded discs (n=5 discs), normalized to background fluorescence of the disc and graphed as a percent of the maximum signal detected. Individual measurement points are shown (light gray), splines created by LOWESS analysis using 10 points per window. (**H-I**) Discs bearing the DR^WNT^ reporter (green) ablated by DC^hepCA^ and expressing *UAS-mir(RHG)* to block cell death, imaged at 36 h AHS. (H) JNK in the absence of cell death activates Wg (red) overlapping the DR^WNT^ reporter (green). (**I**) Mmp1 (red) is induced alongside DR^WNT^ in the absence of Dcp-1 (white), confirming activation is cell-death-independent. Scale bars in all panels = 50 μm, except (F-F’’’’) = 15 μm.

*DR^WNT^-GFP* is activated by JNK signaling (8), likely directly via AP-1 sites since their removal blocks its activity (8). To better characterize this, we monitored *DR^WNT^-GFP* alongside a JNK reporter (*AP-1-RFP*) from 0 h to 36 h AHS (Fig 1E). GFP and RFP are first co-expressed at ∼6 h AHS and largely overlapped thereafter (Fig 1E, 6 h arrowhead). By 12 h AHS, GFP and/or RFP-expressing cells that are Dcp-1–positive are observed, representing dying cells undergoing clearance, while Dcp-1–negative cells indicate those activating the regeneration program (Fig 1F–F’ and S1A–C, arrowheads). This separation of dying and regenerating cells is also observed using a *lexAop-RFP* transgene to label cells undergoing ablation (S1D Fig). At 18 h - 36 h, while a significant overlap of GFP and RFP exists (Fig 1E and S1B Fig), many cells with strong DR^WNT^ enhancer activity showed only weak JNK activity (S1B Fig, arrowheads). A nuclear localized DR^WNT^ reporter (*DR^WNT^[NLS]*) confirms this, with strong enhancer activity in the wound periphery despite low JNK (Fig 1F-F’’’’, arrowheads in F’’’ and F’’’’). Quantification demonstrates a separation of high JNK in the wound center and lower JNK in the periphery (Fig 1F’’’’ and G). Although we cannot rule out a difference in fluorophore perdurance of these reporters, the expression of the known JNK target *Mmp1* mirrors this pattern (S1E Fig), suggesting that JNK alone may not fully explain DR^WNT^ enhancer activity. Since DR^WNT^ represents only the damage-responsive portion of the full DRMS^WNT^ enhancer, we also examined a reporter for the entire DRMS region (*DRMS^WNT^-GFP*, S1F Fig). It showed the same colocalization with JNK-positive cells, with the only difference being weaker GFP signal due to the basal silencing by the MS region (8). Thus, the full length DRMS enhancer behaves similarly to the DR module in its overlap with JNK.

Importantly, developmental JNK signaling (e.g. acting during dorsal closure in the embryo or thorax closure of notum (23–26) fails to activate the DR^WNT^ enhancer even where JNK reporters and targets like *puc* are active (S1G-H Fig). Other JNK-responsive genes like *Mmp1* (27), *Ilp8* (28), and JAK/STAT targets (29) are also not expressed (S1I Fig). A number of these factors are associated with DRMS enhancers (16). These findings imply that additional damage-induced factors are required, or that JNK alone is sufficient but the level of JNK activity dictates the activation of developmental versus damage-specific targets.

To test if JNK alone is sufficient to activate DR^WNT^, we induced JNK signaling while blocking apoptosis to avoid downstream effects of cell death. Apoptotic cells can release extracellular mitogens like Wg and Dpp (30, 31), and generate ROS, triggering JNK in neighboring cells and apoptosis-induced proliferation (AiP) (32–36). To minimize these effects, we used DC^hepCA^ to activate JNK and co-expressed (*UAS-mir(RHG)* (37) to inhibit *rpr*, *hid* and *grim*, blocking apoptosis downstream of AP-1 but upstream of caspase activation. This prevents the signaling events associated with AiP, “undead” cells and feedforward JNK signaling (30, 38, 39), including the generation of ROS via Dronc/Duox (32, 33, 40), which is significantly reduced although not eliminated in this background (S6D Fig). Under these conditions, *DR^WNT^-GFP* can still be activated (Fig 1H), as well as the enhancer’s downstream target *wg* (Fig 1H, arrowhead) and other targets regulated by DR enhancers like *Mmp1* (Fig 1I). This shows that JNK and its immediate downstream effectors are sufficient to trigger the regeneration program via DR enhancers, while cell death and associated signaling events are not required. Interestingly, although blocking cell death in this way prevents undead cell formation and AiP, some ectopic growth still occurs, resulting in larger overall pouch tissue (S1J-K Fig). We speculate that this could be due to the initial activation of growth-promoting factors like Wg (Fig 1H), while the inability to sustain feedforward JNK signaling may limit this growth and prevent the formation of neoplastic tumors observed when apoptosis is blocked at the level of caspase activity (39, 41, 42).

Together, these findings show that JNK can activate DR^WNT^ independently of apoptosis. Moreover, the incomplete overlap between JNK and DR^WNT^ activity, and their distinct developmental expression patterns, suggest additional inputs may modulate enhancer activation, though they likely lie outside cell-death induced signaling events.

### Transcriptomic analysis suggests JAK/STAT can regulate DR enhancers

JNK activates many pathways during regeneration (43), any of which could contribute to DR enhancer regulation. To identify potential candidate pathways, we used next gen sequencing approaches. Previously, we performed ATAC-seq on damaged and undamaged early and late L3 discs to identify genomic regions with altered chromatin accessibility, thus potentially representing DR or DRMS enhancers (Fig 2A) (16). This analysis revealed 243 unique regions; 222 regions in early L3, and 33 in late L3, with increased damage-induced accessibility (16). We re-analyzed these regions for enriched transcription factor (TF) binding motifs that might indicate regulatory signaling pathways (see Materials and Methods). Motif analysis of only early L3 DR regions (Fig 2B), and of all DR regions in early and late L3 discs combined (S2A Fig), revealed enrichment for the chromatin modifier *Trithorax-like* (*Trl*), the DNA binding protein *zeste* (z), which is involved in chromatin-targeted regulation of gene expression, and two early growth response (EGR) family factors, *klumpfuss* (*klu*) and *stripe* (*sr*). The enrichment of these two factors is notable, since previous work in the highly regenerative acoel worm *Hofstenia miamia* found that Egr functions as a pioneer factor to directly regulate early wound-induced genes and activate the entire gene regulatory network necessary for whole body regeneration (44). AP-1 (*kay*/*jra*) motifs were also enriched (Fig 2B and S2A Fig), but at a lower significance level than expected for a known regulator of DR enhancers (p.adj = 3.59x10^-3^ in early L3 DR regions, p.adj = 1.82x10^-5^ in all DR regions), suggesting this analysis can identify factors regulating DR enhancer chromatin accessibility, but may not be sensitive enough to identify TFs controlling their activity.

**Figure 2:**
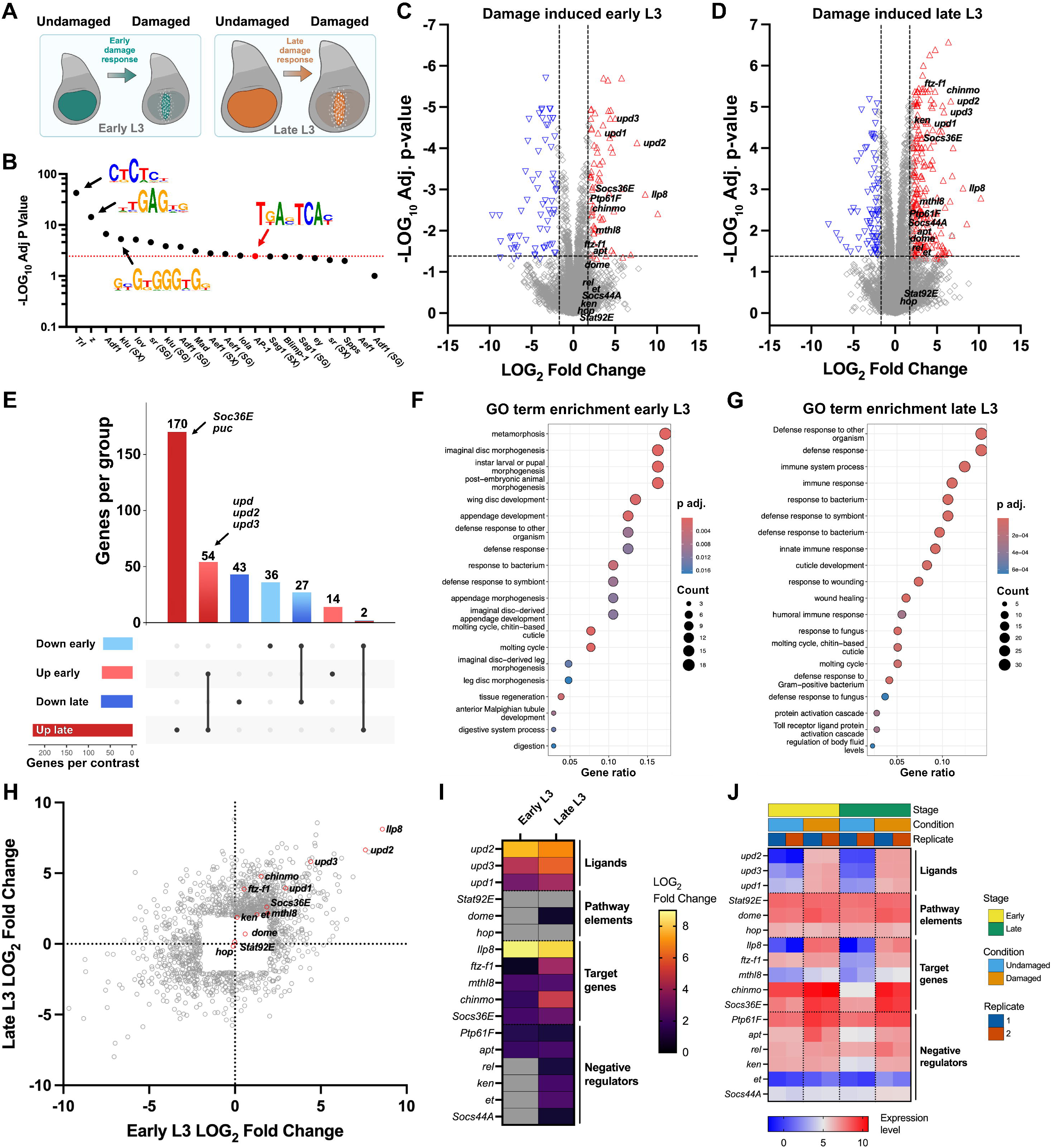
Transcriptomics and motif enrichment identify JAK/STAT as a putative regulator of DR enhancers. (**A**) Schematic of RNA-seq and ATAC-seq (16) conditions used to identify damage-induced changes in early and late third instar (L3) wing discs. Discs were ablated using *rn^ts^>egr* (*rn-GAL4, GAL80ts, UAS-egr*) for 40 hours following culture at 18D°C for 168 h (early) or 216 h (late) after egg deposition (AED), and processed immediately after ablation. (**B**) Motif enrichment analysis (AME) of 222 DR peaks identified in early L3 discs using 7456 static peaks as background. Enriched motifs include binding sites for chromatin regulators *Trl*, *z*, *klu*, and JNK pathway factors *kay/jra* (AP-1). (**C-D**) Volcano plots showing differentially expressed genes (DEGs) in response to damage in (C) early and (D) late L3 discs. Genes are plotted by log_2_ fold change (LOG_2_FC) and -log10 adjusted p-value. JAK/STAT pathway-related genes are highlighted. (**E**) Upset plot showing overlap of differentially expressed genes (DEGs) between early and late L3 stages and damage conditions. Horizontal bars indicate the number of DEGs in each group; vertical bars show shared genes across conditions. *upd* ligands are upregulated in both early and late damaged discs, while *puc* and *Socs36E*, negative regulators of JNK and JAK/STAT, are upregulated only in late L3. (**F-G**) Gene ontology (GO) enrichment among DEGs in early (F) and late (G) L3 damaged discs. The x-axis shows gene ratio; dot size reflects the number of genes in each category, and color indicates adjusted p-value. (**H**) Comparison of LOG_2_FC values between early and late L3 stages for all genes. JAK/STAT pathway genes are labeled in red, others in grey. (**I**) Heatmap showing damage-induced LOG_2_FC in JAK/STAT pathway components, including ligands, signal transducers, targets, and negative regulators. Grey indicates non-significant changes (p > 0.05). (**J**) Heatmap of JAK/STAT pathway gene expression across developmental stages (early vs. late), damage conditions (undamaged vs. damaged), and biological replicates. Red indicates higher expression, blue indicates lower.

To complement this approach, we performed RNA-seq on discs under the same conditions as the ATAC-seq experiment (Fig 2A) to identify genes that are differentially expressed upon damage. In early L3 discs 570 genes were altered (338 upregulated, 232 downregulated, p<0.05, Log_2_FC>0.4, Fig 2C, S1 table), while late L3 discs have 1406 differentially expressed genes (839 upregulated and 567 downregulated, Fig 2D, S1 table). Gene ontology analysis identifies enrichment for transcriptional changes related to imaginal disc morphogenesis, development and tissue regeneration in early L3 discs, and immunity, wound healing and defense responses in late L3 discs (Fig 2F-G). Comparing early and late L3 regenerating discs identifies both stage-specific and shared damage-induced genes, (Fig 2E, S2 table), including all three *upd* ligands of the JAK/STAT pathway upregulated at both stages (Fig 2E). JAK/STAT signaling is known to function during stress responses (28, 29, 45–49), being activated downstream of JNK in these contexts (22, 28, 29, 47, 50, 51). While the expression of the JAK/STAT core pathway components, *domeless* (*dome*), *hopscotch* (*hop*) and *Stat92E* remain largely unchanged in our RNA-seq data (Fig 2H-J), several known targets such as *Ilp8* (28), *ftz-F1* (52), *mthl8* (53), *chinmo* (54) and *Soc36E* (55) are upregulated by damage in both early and late L3 discs, suggesting pathway activation and thus the potential to regulate DR enhancers. This specific transcriptomic behavior of ligands, pathway components and target genes mirrors that of the known DR enhancer regulator, JNK (S2B-E Fig). Furthermore, although the binding motif for Stat92E (56) is not globally enriched in the DR regions versus static peaks identified by ATAC-seq (Fig 2B and S2A Fig), a targeted search found 50.62% (123/243) of DR regions contain Stat92E binding sites, compared to 68.72% (167/243) with AP-1 sites (FIMO tool p<0.001, meme-suite.org). This aligns with published single cell ATAC-seq data showing AP-1 and Stat92E motif enrichment in chromatin regions activated in wound responsive cells (49). Notably, this enrichment occurs within a specific subset of cells within the wound, which may explain its absence from the analyses of our data that used whole discs (16). Together, these findings suggest JAK/STAT signaling may act alongside JNK to regulate DR enhancers.

Interestingly, despite damage-induced *10XSTAT-GFP* reporter activity being diminished in ablated late L3 discs (16), our RNA-seq data showed robust *upd* ligand expression. However, it also shows upregulation of several negative regulators of JAK/STAT (Fig 2I-J), such as *apontic* (*apt*), which is known to restrict pathway activity (16, 57). This suggests that while JAK/STAT signaling is still induced in late L3, its output might be limited by stage-specific negative regulators.

### JAK/STAT signaling coincides with DR enhancer activity

To test whether JAK/STAT regulates DR enhancers, we examined the spatial and temporal dynamics of JAK/STAT and JNK signaling using a fast-turnover GFP reporter for JAK/STAT (*10XSTAT-DGFP*) (58) and *AP-1-RFP* as before (59). Prior to injury, JAK/STAT is active in the hinge while JNK is absent (Fig 3A). Following DC^hepCA^ ablation, JNK is detected by ∼6 h and coincides with DR^WNT^ (Fig 1E). However, JAK/STAT is not detected in the pouch until 12 h AHS (S3A Fig), consistent with its activation downstream of JNK. By 18 h AHS, JNK can be distinguished in regions of higher (distal) and lower (proximal) activity (Fig 3B and 3D, S3A Fig), while JAK/STAT appears across both regions (Fig 3B). At 24 h AHS, JNK levels remains elevated distally (Fig 3C, S3A), but JAK/STAT activity declines particularly in the distal pouch (Fig 3C-D, S3A Fig). Quantification shows these patterns diverge over time (Fig 3F-G), with JNK remaining centrally within the wound, and JAK/STAT localizing more peripherally, until both pathways decline by 36 h AHS (Fig 3F-G and S3A-B Fig). These findings agree with recently published work that used a different genetic ablation system (*rn-GAL4/GAL80ts/UAS-egr*) to demonstrate that JNK and JAK/STAT are activated in response to injury and subsequently resolve into separate regions due to a mutually repressive interaction (48). This separation allows JNK-positive cells to transiently pause proliferation, while JAK/STAT promotes it in surrounding cells (48). This separation is also observed via single cell analyses (49). Using DC^hepCA^, we observed a similar pause in proliferation in the high JNK cells at 18 h and 24 h AHS, which is relieved by 36 h AHS (Fig 3H-J), while the surrounding lower JNK cells do not show this pause (Fig 3H-J). Thus, despite the different timing of ablation and recovery imposed by each ablation method, a similar pattern of wound signaling and proliferation is observed, while the acute injury caused by the DC system further reveals that cells have distinct responses based on JNK levels.

**Figure 3:**
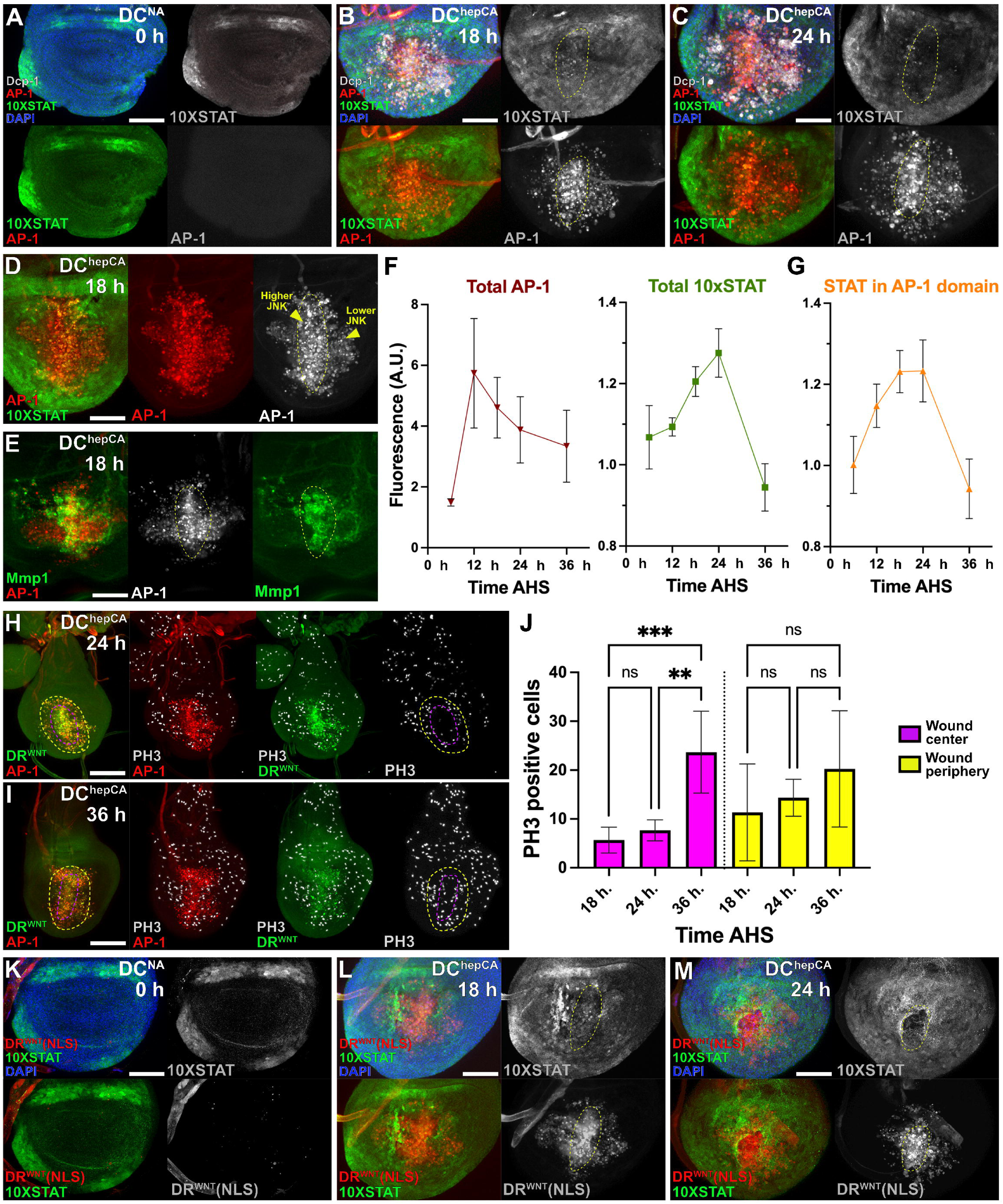
DR^WNT^ overlaps with dynamic changes in JNK and JAK/STAT expression during regeneration. (**A-C**) Time course of discs ablated with DC^hepCA^, imaged at 0, 18, and 24 h after heat shock (AHS), showing JNK activity (*AP-1-RFP*, red), JAK/STAT activity (*10XSTAT-DGFP*, green), and cell death (Dcp-1, white). At 0Dh, only developmental reporter expression is seen, with JAK/STAT restricted to the hinge. At 18Dh, JNK is strongly induced in the wound center (dotted circle), and JAK/STAT is upregulated both within the wound and non-autonomously in surrounding tissue. By 24Dh, JAK/STAT is reduced in the JNK-high wound center, while JNK remains elevated. (**D**) At 18Dh AHS, discrete high and low JNK zones (red, dotted outlines) are visible in the distal pouch, coincident with JAK/STAT activity (yellow arrowheads). (**E**) Expression of JNK target Mmp1 (green) at 18 h highlights spatially distinct regions of high vs. low JNK at 18Dh AHS (dotted outline). (**F**) Quantification of JNK (AP-1-RFP) and JAK/STAT (10XSTAT-DGFP) reporter intensity in the pouch, normalized to background, from 6– 36Dh AHS (n = 21 discs; error bars are SD; arbitrary units (A.U.) (**G**) Quantification of JAK/STAT activity specifically within the JNK-positive domain (as in F), showing early co-activation followed by reduction of JAK/STAT signal in JNK-high regions and continued increase in peripheral regions (n = 21; error bars are SD). (**H-I**) Discs imaged at 24Dh (H) and 36Dh (I) AHS stained for proliferation (PH3, white), with JNK (red) and DR^WNT^ (green). At 24Dh, high JNK regions (purple dashed outline) lack PH3+ cells, while JNK-low periphery (yellow outline) shows ongoing proliferation. At 36Dh, proliferation is restored in the JNK-high zone. (**J**) Quantification of PH3+ cells within high and low JNK regions from 18–36Dh AHS. High JNK regions show significantly reduced proliferation at 18Dh and 24Dh, recovering by 36Dh (ordinary one-way ANOVA: 18Dh p = 0.006, 24Dh *p = 0.0004, 36Dh p > 0.05; n = 23 discs; error bars are SD). Proliferation in low JNK areas increases over time but differences are not statistically significant. **(K-M)** Discs expressing DR^WNT^[NLS] (red) and *10XSTAT-DGFP* (green), either unablated (DC^NA^, K) or ablated (DC^hepCA^, L–M). At 0Dh AHS (K), no DR^WNT^ activity is observed, and JAK/STAT is limited to the hinge. At 18Dh (L), DR^WNT^ overlaps with JAK/STAT in both the wound center and periphery. At 24Dh (M), JAK/STAT decreases in the center while DR^WNT^ remains active (yellow dashed outlines). Scale bars in all panels = 50 μm, except (A-C) and (K-M) = 15 μm.

We next analyzed *DR^WNT^-GFP* enhancer activity in this dynamic context (Fig 3K-M). Early *DR^WNT^-GFP* activity (0 h to 12 h AHS) significantly overlaps with JNK (Fig 1E), then expands by 18 h AHS to include peripheral cells with lower JNK and new JAK/STAT activity (Fig 1F–F’’’’, 3B, 3L). At 24 h AHS, DR^WNT^ remains active in both central high JNK and peripheral JAK/STAT-positive regions, even as central JAK/STAT declines (Fig 3M), likely due to repression by JNK (48). Together, these results show that the DR^WNT^ enhancer activation begins in JNK-positive cells and later spreads to include JAK/STAT positive regions in the wound periphery. This dynamic overlap suggests that both pathways may contribute to DR^WNT^ activity in a developing wound.

### JAK/STAT directly regulates DR enhancer activity in the wound periphery

To test whether JAK/STAT is required for DR enhancer activation, we knocked down *Stat92E* (*UAS-Stat92E^RNAi^*) during regeneration and assessed *DR^WNT^-GFP*. *Stat92E* knock down significantly reduced *DR^WNT^-GFP* activity throughout regeneration (Fig 4A-B), as well as activity of other DR enhancers including *DRMS^WNT^-GFP* and *DR^Mmp1^-GFP* (S4A–G Fig). Importantly, this reduction is most apparent in the wound periphery (Fig 4A’), where JNK levels are low and JAK/STAT is normally active (Fig 1F, 3B– D, 3L), indicating that JAK/STAT is necessary for enhancer activation in this domain. *Stat92E* knockdown does not affect cell death (Fig 4C-C’) or JNK signaling (Fig 4D-D’), placing JAK/STAT function downstream or parallel to JNK. To assess the spatial requirement for JAK/STAT in the center of the wound versus the periphery, we knocked down *Stat92E* specifically in the posterior compartment using *hedgehog*-GAL4 (*hh-GAL4*), thus affecting only half the wound (Fig 4E). DR^WNT^ activity was mostly preserved centrally but was reduced in the posterior wound periphery (Fig 4E), reinforcing the possibility that JAK/STAT is required for DR^WNT^ activity at the wound periphery, where JNK is weaker.

**Figure 4:**
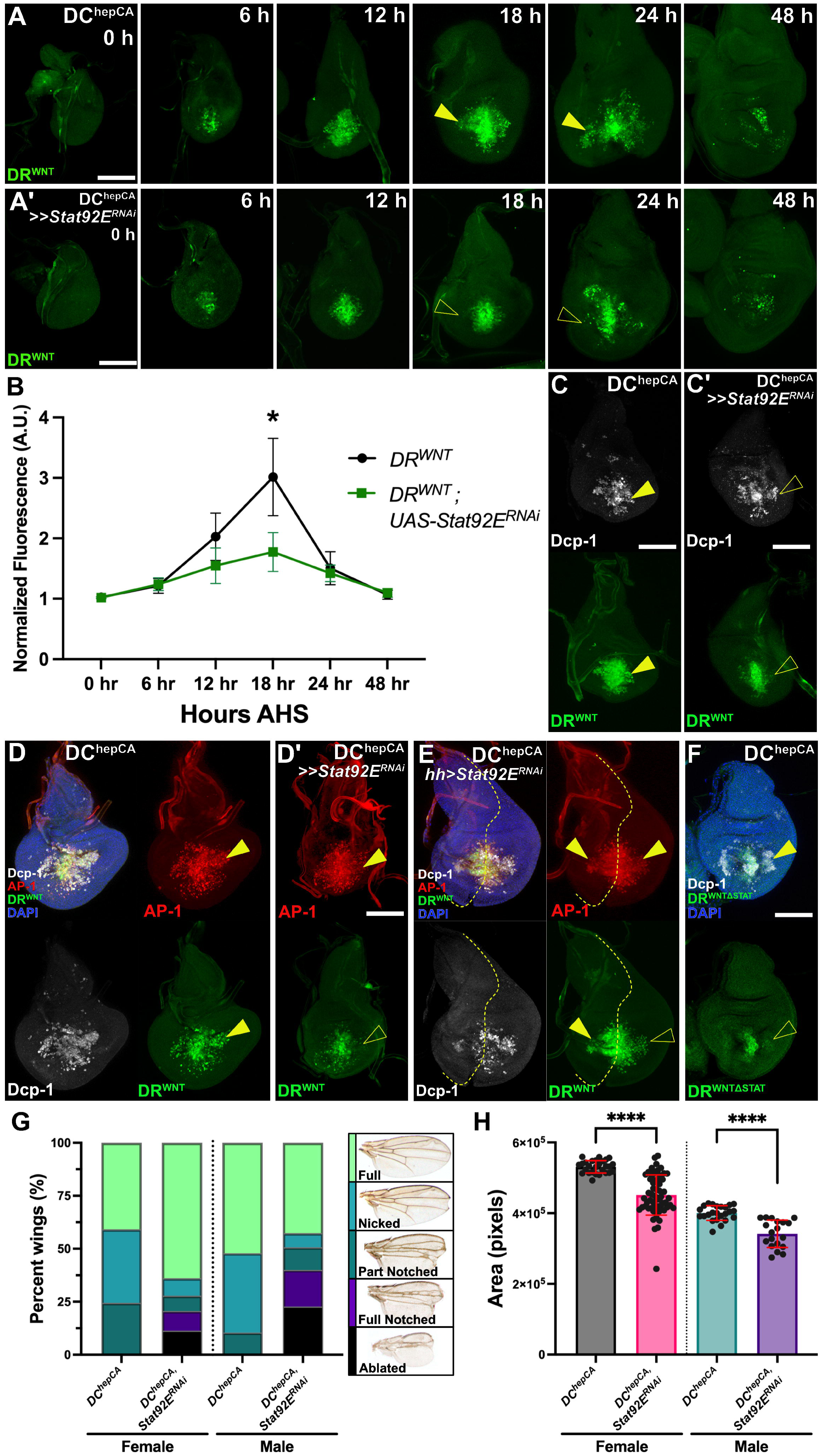
Stat92E is necessary for full activation and lateral domain activity of DR^WNT^. (**A-A’**) Time course (0–48 h AHS) of *DR^WNT^-GFP* (green) reporter activation in DC^hepCA^-ablated discs with (A) or without (A’) *Stat92E* knockdown. Peripheral DR^WNT^ activity is reduced at 18 and 24 h AHS with *Stat92E^RNAi^* (arrowheads). (**B**) Quantification of DR^WNT^ fluorescence (normalized to background) confirms significant reduction at 18 h AHS with *Stat92E^RNAi^*. Unpaired t-tests: *p < 0.000001; n = 44 (control), n = 45 (*Stat92E^RNAi^*). A.U. = Arbitrary Units, (**C-C’**) *DR^WNT^-GFP* (green) and Dcp-1 (white) at 18 h AHS show similar cell death with or without *Stat92E*, but loss of peripheral DR^WNT^ (yellow arrowheads) in knockdown. (**D-D’**) Discs as in (C-C’) with the JNK reporter (AP-1, red). The peripheral DR^WNT^ activity (yellow arrowhead) is lost upon *Stat92E* knockdown, while JNK activity is unaffected. (**E**) Posterior-specific *Stat92E* knockdown (*hh-GAL4*) shows reduced DR^WNT^ in posterior wound periphery (yellow arrowheads) compared to wild-type anterior; A/P boundary indicated by dashed line. (**F**) *DR^WNT^*^Δ*STAT*^*-GFP* (lacking *Stat92E* sites) shows loss of peripheral activity at 18 h AHS (yellow arrowhead). (**G**) Adult wing phenotype scoring post-ablation shows impaired regeneration with *Stat92E^RNAi^*; n = 127 (control), n = 172 (*Stat92E^RNAi^*), separated by sex. (**H**) Wing size is reduced in adults with *Stat92E^RNAi^*; unpaired t-tests: ****p < 0.0001; n = 51 (control), n = 78 (*Stat92E^RNAi^*), data by sex. Scale bars in all panels = 50 μm.

The behavior of another DRMS enhancer identified in our accessibility data set (16) is consistent with this idea. A GFP reporter for DRMS^TDC1/2^, an enhancer located near to the *Tdc1* and *Tdc2* genes (S4H Fig), which are both highly upregulated upon damage (S1 Table), contains AP-1 but not Stat92E consensus sites (S4H Fig), and exhibits damage-responsive, maturity-silenced behavior (16). *DRMS^TDC1/2^-GFP* shows no activity in the periphery of a wound and is unaffected by *Stat92E* knockdown (S4I–J Fig), suggesting it is JNK-dependent but JAK/STAT-independent. In this experiment, the minimal DR module for this enhancer has not been identified, therefore the full DRMS is used To test whether JAK/STAT acts directly on DR^WNT^, we generated a mutant DR^WNT^ reporter lacking its three identified consensus Stat92E binding sites (*DR^WNT^*^Δ*STAT*^*-GFP*, S4K Fig). This reporter showed reduced activity in the wound periphery (Fig 4F and S4L Fig), phenocopying the *Stat92E* knockdown (Fig 4A-D’), suggesting JAK/STAT regulates the enhancer directly.

As noted, JAK/STAT is well established as necessary for regeneration (28, 29, 45–49), and indeed the knockdown of *Stat92E* following ablation by DC^hepCA^ strongly limits regeneration in this system, as assayed by both wing scoring (Fig 4G) and wing area measurements (Fig 4H). Although JAK/STAT supports regeneration through pleiotropic effects, including promoting cell survival and proliferation (45), these findings highlight an additional mechanism through which JAK/STAT could promote regeneration: by directly regulating the activity of DR enhancers and thus their targets.

### Ectopic JAK/STAT is insufficient to activate DR enhancers

Although JAK/STAT signaling is necessary for the full spatial activity of DR enhancers downstream of JNK, the question remains as to whether it is sufficient to induce their activity in the absence of JNK. To test this, we overexpressed *hop* in the distal pouch (*salm-GAL4*, *UAS-hop48a*) to ectopically activate JAK/STAT signaling. Although this expression activates *10XSTAT-DGFP* in the pouch (Fig 5A), it is insufficient to activate the *DR^WNT^-GFP* reporter (Fig 5B). This is even despite the presence of low levels of cell death caused by *hop* overexpression (Fig 5A-B). We repeated this experiment with different GAL4 drivers that express throughout the whole pouch (*rotund-GAL4*) or in both the pouch and the notum (*hh-GAL4 and ptc-GAL4*). These drivers activate JAK/STAT signaling (S5A Fig and S5C Fig) and even can lead to the formation of ectopic pouch tissue (S5C-D Fig) as described previously (51). However, they do not activate the DR^WNT^ enhancer (S5B Fig and S5D Fig) or other JNK targets such as *Mmp1* (S5A-B Fig). The use of a stronger *hop* construct (*UAS-hopORF*) leads to significant cell death, which in turn triggers JNK signaling shown by *Mmp1*, and thus JAK/STAT and *DR^WNT^-GFP* expression (S5E-F Fig). These data show that JAK/STAT signaling alone is not sufficient to activate DR enhancers, but rather is required to fully promote and maintain their activity after activation by JNK.

**Figure 5:**
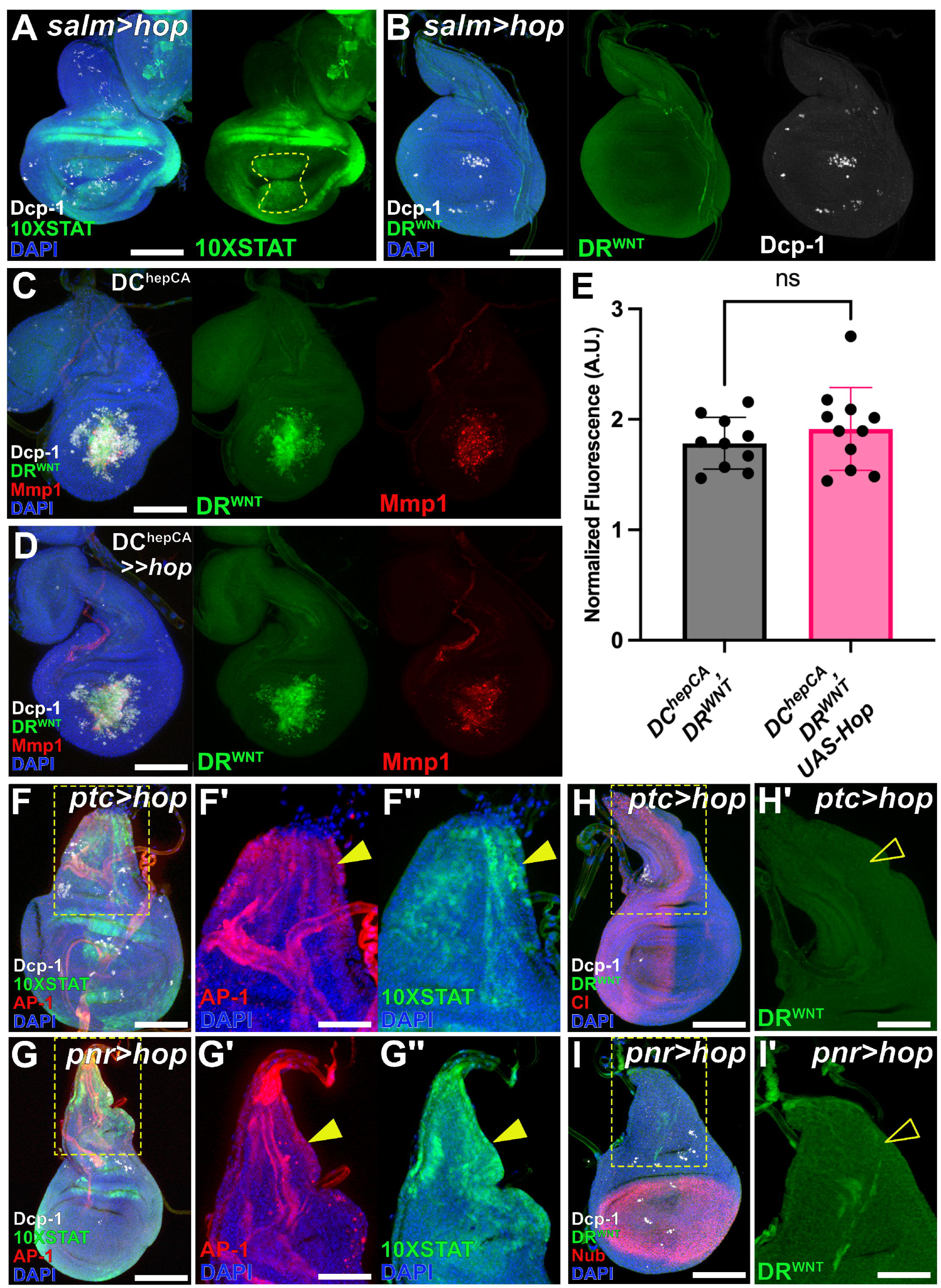
JAK/STAT activation is not sufficient to induce DR^WNT^. (**A-B**) Ectopic JAK/STAT signaling in the distal pouch (*salm-GAL4*, *UAS-hop*; yellow outline) activates *10XSTAT-DGFP* (green in A) without inducing cell death (Dcp-1, white) or DR^WNT^ reporter expression (green in B). (**C–D**) In DC^hepCA^-ablated discs, DR^WNT^ is activated (green) alongside Dcp-1 (white) and Mmp1 (red, JNK activity). Ectopic JAK/STAT (*UAS-hop*) does not alter DR^WNT^ or JNK activity. (**E**) Quantification of DR^WNT^ reporter intensity shows no significant difference with ectopic JAK/STAT. n = 10 (*DC^hepCA^*), n = 11 (*DC^hepCA^, UAS-hop*); unpaired two-tailed t-test, ns (not significant). A.U. = Arbitrary Units. (**F-F’’**) Overexpression of *hop* in an A/P stripe (*ptc-GAL4, UAS-hop*) increases *10XSTAT-DGFP* (green) and JNK (AP-1-RFP, red) in the notum (arrowheads), without cell death (Dcp-1, white). Insets (F’–F’’) show magnified views. (**G-G’’**) Ectopic JAK/STAT in the notum (*pnr-GAL4, UAS-hop*) similarly elevates STAT and JNK reporters (green and red), without cell death. (**H-H’**) DR^WNT^ reporter (green) is not activated by A/P stripe JAK/STAT (open arrowhead in H’); Ci (red) marks the A/P boundary, Dcp-1 (white) shows no cell death. (**I**) Ectopic JAK/STAT in the notum (*pnr-GAL4, UAS-hop*) also fails to activate DR^WNT^ (green; open arrowhead in I’). Nub (red) marks the pouch, Dcp-1 (white) labels apoptosis. Scale bars in all panels = 50 μm, except (F’-F’’,G’-G’’,H’,I’) = 25 μm.

Since JNK is a prerequisite for DR enhancer activation, we wondered whether JAK/STAT could alter DR enhancer behavior in different contexts where JNK is already present: either following damage or during normal development. We first tested ectopically expressing *hop* during ablation and found that additional JAK/STAT signaling does not significantly alter *DR^WNT^-GFP* expression (Fig 5C-E), suggesting that JAK/STAT is not limiting for DR enhancer activity during regeneration. Next, to examine whether JNK that occurs during development can activate *DR^WNT^-GFP* when JAK/STAT is introduced, we expressed *hop* in the notum, where endogenous JNK promotes thorax closure and JAK/STAT is normally absent (23, 24, 60) (S1I Fig). We used two different tissue-specific GAL4 drivers, *patched* and *pannier*, (*ptc-GAL4* and *pnr-GAL4*), both of which express in domains that overlap JNK in the notum (Fig 5F-G’’ and S5G-H Fig) and activate the 10XSTAT reporter in response to *hop* (Fig 5F’’ and G’’). Even with the activity of both signaling pathways, the *DR^WNT^-GFP* reporter failed to activate (Fig 5H-I’), suggesting that JNK expressed during development is not sufficient to activate DR enhancers, even with the addition of JAK/STAT. By contrast, JNK during injury initiates DR enhancer activity, while the subsequent induction of JAK/STAT allows their expansion into the wound periphery. Since cell death and its associated downstream signaling are dispensable for DR enhancer activity (Fig 1), this reduces the likelihood that injury-specific factors are responsible for this context-dependent activation and raises the possibility that JNK signaling intensity might instead regulate DR enhancer activation.

### A threshold level of JNK activity is required to initiate the regeneration program via DR enhancers

To test whether the level of JNK signaling influences DR enhancer activation, we developed a method to induce different intensities of steady-state JNK activity, unlike the dynamic changes that occur within a wound. Using the temperature-sensitivity of GAL4 (61), we expressed a wild type *hep* transgene in the distal pouch (*salm-GAL4, UAS-hep^WT^*) at 22°C for JNK^Low^ and 30°C for JNK^Med^, and a constitutively active *hep* (*salm-GAL4, UAS-hep^CA,^ GAL80^ts^*) activated by temporary culture at 30°C for JNK^High^. Cell death was blocked with *mir(RHG)* to reduce ROS and limit feedforward JNK (22, 40, 62). Indeed, ROS is only minimally produced under these conditions versus ablation (S6C-C’’ Fig), and JNK activity is consequently confined to the distal pouch (Fig 6C). Expression of the JNK target *mol* that is important for ROS production is also unchanged (S6E-F’’ Fig). We confirmed different JNK activity levels using *AP-1-RFP* quantified via fluorescence intensity (Fig 6A) and qRT-PCR of transcript expression (Fig 6B).

**Figure 6:**
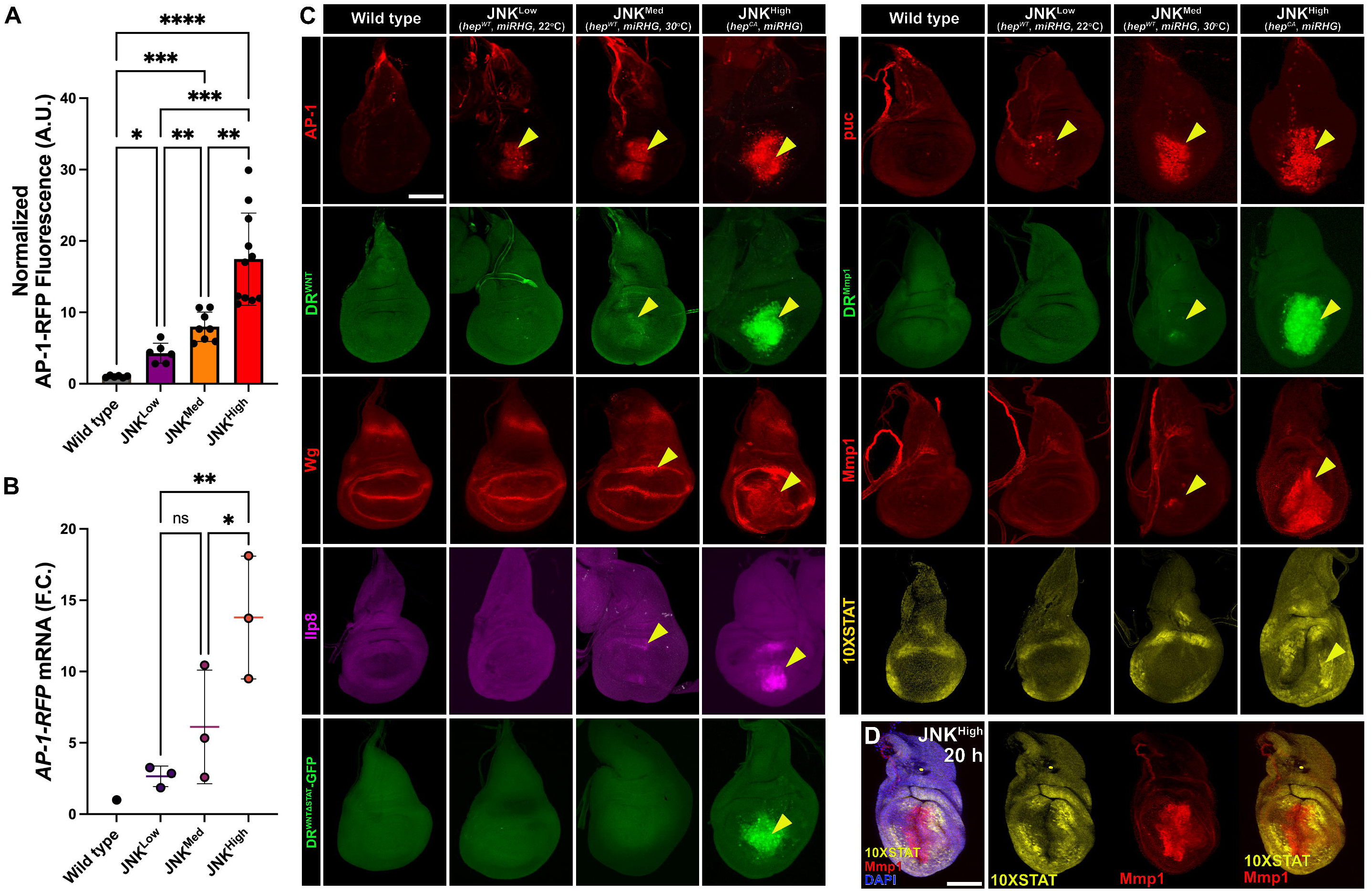
A threshold level of JNK signaling is required to activate DR^WNT^ and downstream targets. **(A)** Quantification of JNK levels (AP-1-RFP fluorescence, normalized to background) in JNK^Low^ (salm > hep^WT^, 22°C), JNK^Med^ (salm > hep^WT^, 30°C), and JNK^High^ (salm > hep^CA^) discs, all with blocked apoptosis (*UAS-mir[RHG])*. One-way ANOVA: *p = 0.0108 (WT– JNK^Low^), ***p = 0.0002 (WT– JNK^Med^), ****p < 0.0001 (WT– JNK^High^), **p = 0.0097 (JNK^Low^ – JNK^Med^), ***p = 0.0002 (JNK^Low^ – JNK^High^), **p < 0.0031 (JNK^Med^ – JNK^High^). Error bars: SD; n = 6 (WT), 7 (JNK^Low^), 8 (JNK^Med^), 11 (JNK^High^); A.U. = Arbitrary Units., **(B)** Measurement of JNK levels in conditions of JNK^Low^, JNK^Med^, and JNK^High^ (as above) via qRT-PCR of *AP-1-RFP* transcript levels. Graphs show the mean fold change (F.C.) of *AP-1-RFP* in each condition relative to the wild-type (physiological level of *AP-1-RFP* in uninjured discs at the specified temperature). Ordinary one-way ANOVA between JNK conditions, **p=0.0072 (JNK^Low^-JNK^High^), *p<0.033 (JNK^Med^-JNK^High^). Error bars are SD, ns = not significant (JNK^Low^-JNK^Med^), n = 3 independent biological repeats per condition. **(C)** Reporter activation under increasing JNK levels*. AP-1-RFP* and *puc-LacZ* (JNK reporters) activate at all levels (yellow arrowheads). DR^WNT^, DR^Mmp1^, Mmp1, and Ilp8 activate weakly at JNK^Med^ and strongly at JNK^High^. 10XSTAT reporter is activated only at JNK^High^. **(D)** Under JNK^High^ conditions, *10XSTAT-DGFP* (yellow) and the JNK target Mmp1 (red) become spatially distinct, emulating a mature-stage wound. Scale bar in all panels = 50 μm.

With this setup, we assessed DR enhancers and known regeneration targets *Mmp1*, *Ilp8*, *wg*, and *Stat92E* (Fig 6C). JNK^Low^ failed to activate *DR^WNT^-GFP* or JAK/STAT signaling, although it did activate *puc*, which likely has a lower activation threshold. JNK^Med^ induced weak DR^WNT^ and DR^Mmp1^ reporter activity, but not robust expression of other targets. By contrast, JNK^High^ strongly activates DR enhancers, including DR^WNTΔSTAT^, and all tested regeneration genes within the high-JNK domain (Fig 6C). Interestingly, JAK/STAT activation occurred outside this domain (Fig 6D), consistent with the non-autonomous induction observed in wounds (48), but did not lead to DR enhancer activation there, likely because *mir(RHG)* prevents JNK from spreading and forming a low-JNK domain needed for co-activation with JAK/STAT. Thus, these experiments demonstrate three important findings: 1) a threshold level of JNK is needed to activate JAK/STAT signaling, which in steady-state conditions (and a mature wound) becomes exclusively non-autonomous outside of the area of JNK activity, 2) at high levels of JNK, DR enhancers and their downstream pro-regeneration targets can be activated in the absence of JAK/STAT, and 3) when the spread of JNK and the subsequent formation of different levels is prevented, DR enhancer activation outside of the high JNK domain does not occur, even in the presence of JAK/STAT.

Finally, to confirm the requirement for lower JNK combined with JAK/STAT to promote normal DR enhancer behavior in a wound context, we observed manipulations of these factors in DC^hepCA^ ablation (Fig 7A-A’’’). Knockdown of *Stat92E^RNAi^* inhibits the spread of DR enhancer activity into the wound periphery where JNK is lower (Fig 7B-B’’’) but does not affect ROS production (S6D’ Fig) or JNK levels (Fig 7B-B’’’). Blocking cell death with *mir(RHG)* to prevent the formation of different JNK levels within the wound restricts *DR^WNT^-GFP* to the high JNK domain (Fig 7C-C’’’). Combining *mir(RHG)* and *Stat92E^RNAi^* phenocopies this result (Fig 7D-D’’’). Together, these results show that DR enhancer activation during regeneration requires either high JNK at the wound center or the combination of low JNK and JAK/STAT in the periphery.

**Figure 7:**
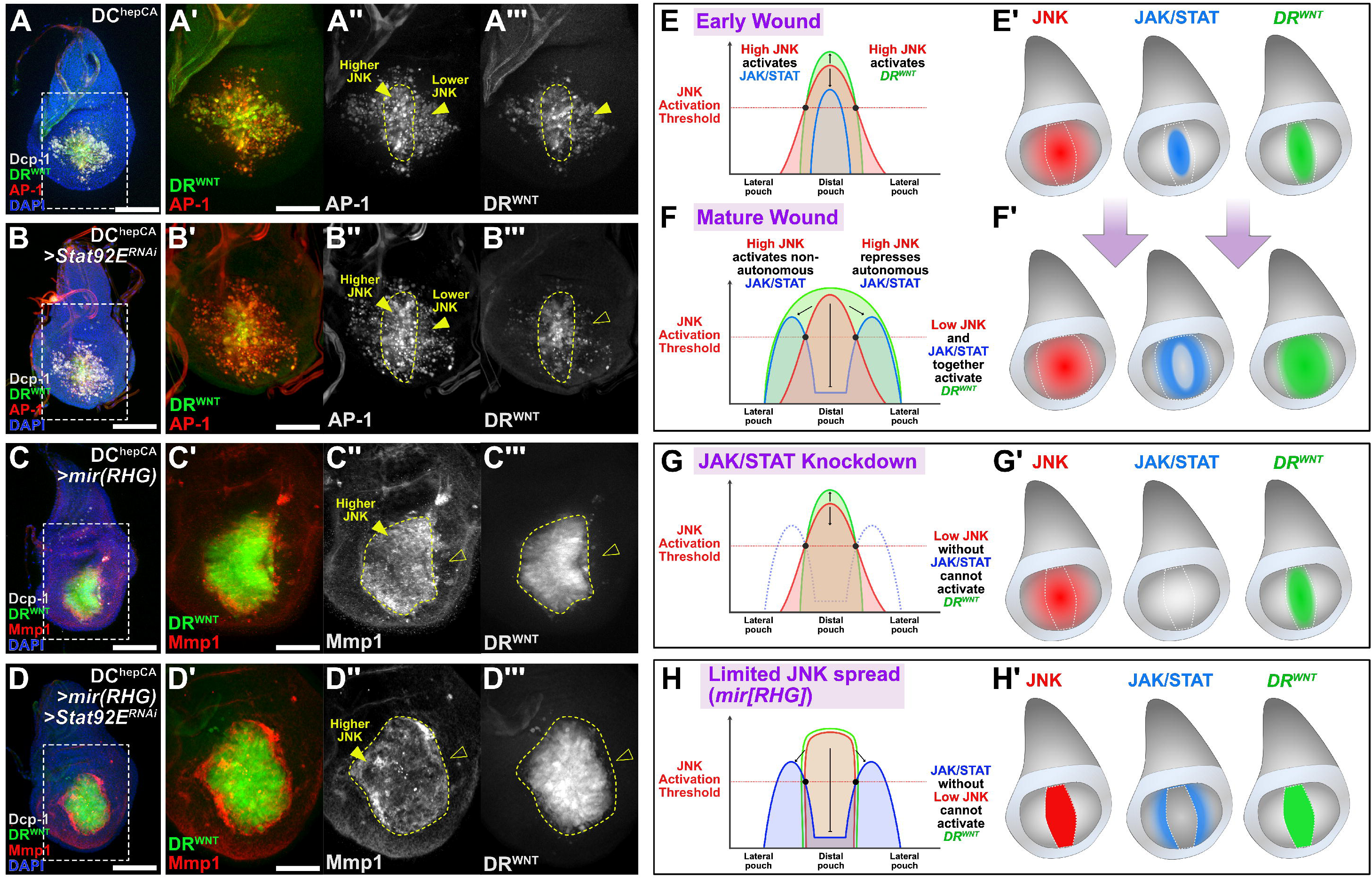
JNK and JAK/STAT expression dynamically activates DR^WNT^ during wound formation. (**A-D’’’**) Discs bearing the DR^WNT^ reporter (green) and the AP-1 reporter (red in A and B) or stained for JNK target Mmp1 (red in C and D), ablated by DC^HepCA^. Discs are wild type (A), with *Stat92E* knockdown (B), with cell death blocked by *mir(RHG)* (C) or with *Stat92E* and cell death block by *mir(RHG)* together (D). DR^WNT^ activity occurs in cells of both high and low JNK of a wound (A-A’), while lack of JAK/STAT limits DR^WNT^ activity to cells with high JNK (B-B’’’). Blocking cell death via *mir(RHG)* to inhibit ROS-mediated spread of JNK prevents the formation of a low JNK area where JAK/STAT can be activated, and therefore DR^WNT^ is limited to the high JNK cells in the center of the distal pouch (C-C’’’). Preventing the spread of JNK via blocking cell death with *mir(RHG)* and blocking JAK/STAT shows the same lack of DR^WNT^ activity outside of the distal pouch (D-D’’’). Limited JNK spread is shown by Mmp1 staining in C and D. Yellow dashed outlines indicate area of high JNK. Yellow arrowheads indicate high JNK cells in the center of the wound and lower JNK cells in the wound periphery. Open arrowheads indicate lack of DR^WNT^ activity in B’’’, C’’’ and D’’’’, and lack of JNK activity shown by Mmp1 in C’’ and D’’. (**E-H’**) Graphical representation of the mechanism by which JNK and JAK/STAT expression leads to DR^WNT^ activity across a developing wound in early L3 larvae from the initial injury (up to ∼18 h AHS) to the mature injury (after ∼18 h AHS), (E-F’), and in conditions of JAK/STAT knockdown (G-G’) or when JNK signaling spread is prevented (H-H’). See main text for details. Scale bars in (A), (B), (C) and (D) = 50 μm, other panels = 25 μm.

Overall, these data support a model in which spatially distinct thresholds of JNK regulate regenerative gene expression (Fig 7E-H’). High JNK in the center of the wound is sufficient to activate DR enhancers (Fig 7E-E’). This high JNK also induces JAK/STAT signaling, which later becomes repressed in these same cells by JNK but activated non-autonomously in adjacent low JNK tissue (Fig 7F-F’). In this peripheral domain the combination of JAK/STAT and low JNK co-activate DR enhancers (Fig 7F-F’).

When JAK/STAT is removed, DR enhancers activity in this region is lost (Fig 7G-G’). Similarly, when cell death is blocked by *mir(RHG)*, limiting the spread of JNK beyond the center of the wound, DR enhancer activity becomes confined to the high JNK domain (Fig H-H’). This mechanism ensures that DR enhancers respond proportionally to injury: high JNK initiates the response, while JAK/STAT extends enhancer activation outward. Since DR enhancers likely regulate diverse genes (16), these JNK thresholds provide a spatial framework for controlling the magnitude, timing, and domain of regenerative gene expression, and distinguishes between the activation of targets required for regeneration versus normal development.

## Discussion

Our previous work identified and characterized DRMS enhancers; regulatory elements within the *Drosophila* genome that activate regeneration-promoting genes in response to damage, but become silenced with maturity to limit tissue repair (8, 16). These enhancers are activated by JNK signaling via a damage-responsive (DR) region, although it is unclear why this activation is injury-specific and is absent from other JNK-dependent contexts like embryogenesis or pupariation (23–26). It is also unknown how a single pathway like JNK produces spatially dynamic gene expression during regeneration.

We have addressed both issues here. Firstly, by isolating JNK signaling from the downstream events of apoptotic signaling, we have shown that DR enhancers activate independently of cell death, requiring only JNK and its immediate targets. One such target, JAK/STAT, is an additional direct input into enhancer activity alongside JNK. We find that high JNK levels are sufficient to initiate DR enhancer activity, while JAK/STAT amplifies and expands this activity into wound-proximal regions where JNK alone is insufficient. Below this threshold, only certain JNK targets like those necessary during development (*puc),* are expressed.

Within a wound, apoptosis promotes ROS-mediated P38 and high JNK activity in surrounding cells (22, 63), activating DR enhancers and their targets (e.g. *wg*,*Mmp1)*, alongside promoting a transient senescence-like state (48). JNK also induces expression of *upd* ligands, activating JAK/STAT signaling (22, 28, 29, 47, 50, 51), which initially acts in an autocrine fashion within the wound, but subsequently becomes paracrine in the wound periphery with low JNK (48). In these outer regions, JAK/STAT and JNK enables DR enhancer activity, thus extending and sustaining regeneration beyond the initial wound. When JAK/STAT is inhibited or JNK is spatially restricted by blocking cell death signaling, peripheral enhancer activation fails, thus limiting the regeneration program to the immediate wound site. Thus, DR enhancers not only initiate regenerative gene expression downstream of JNK but also translate heterogenous levels of JNK into dynamic spatial and temporal expression patterns required for regeneration.

### DRMS enhancers as a mechanism to spatially regulate regenerative gene expression

Regeneration-specific enhancers have been identified in diverse organisms and tissues (5, 6, 64). In highly regenerative Zebrafish, such elements at *leptin B* and *ctgfa* are involved in heart and fin regeneration (9, 65), with numerous other damage-responsive loci found genome-wide (66, 67). Such regulatory elements likely function in other injury models, potentially even in humans (68, 69). Their existence aligns with the need for regeneration-specific expression patterns, those that are unique to wound healing, or that bridge wound healing to developmental programs. Indeed, genome-wide studies in axolotl, Xenopus, mouse and *Drosophila* regeneration confirm such transcriptional and epigenetic profiles (49, 70–74).

Our work in imaginal discs suggests regeneration enhancers do more than initiate reparative gene expression, but also reinforce heterogeneity in the regenerative response. A single pathway, JNK, simultaneously activates multiple genes, enabling a rapid and coordinated response to damage. However, differences in JNK levels caused by proximity to injury or tissue identity (75), must be decoded to produce appropriate spatial gene expression. DR enhancers enable this by translating graded JNK activity into differential target gene expression needed for recovery. Comparisons of full versus partial disc ablation (8, 16), or clonal patches of varying size that are triggered to die (8), supports the idea that DR enhancers drive spatial expression diversity based on JNK intensity. Single cell sequencing of ablated wing discs also corroborates this (49, 74), showing that cell populations with discrete cellular identities form following injury and change over time. Injuries within the wing pouch become striated through the formation of a central JNK-expressing population, which downregulates markers for proliferation and expresses factors to limit differentiation like *Ets21C*, alongside upregulation of paracrine signaling factors like *wg* and the *upds* ligands (48, 49, 74). Cells of the periphery receive these signals, gain JAK/STAT activation and express factors that support survival and proliferation, (48, 49, 74). This regionalization, originally stemming from JNK signaling, is crucial for coordinating growth and repatterning during regeneration (48), facilitated by DR enhancers.

### Signaling factors that activate and regulate DR enhancers are potentially conserved

JNK and JAK/STAT signaling are both essential for regeneration in *Drosophila*, with complex roles in proliferation, cell survival and identity (43, 45)(28, 29, 45–48). In many cases, JAK/STAT in the context of a wound depends on prior JNK activity, raising the possibility that DR enhancers could mediate any events in regeneration that rely on input from both pathways. JNK is closely associated with regeneration enhancers in many if not all known examples, even in organisms with highly contrasting abilities to regenerate like zebrafish and mammals (9, 66, 76–78), and in diverse tissues, from imaginal discs to Schwann cells (8, 16, 18, 79), implying it is likely a conserved input (6). JAK/STAT, though not yet linked to DR enhancers in other species, is essential for regeneration in many contexts. For example, Zebrafish fin regeneration is impaired without *Stat3* due to aberrations in macrophage recruitment (80). Thus, it will be important to test whether JAK/STAT plays a similar direct role in DR enhancer regulation outside *Drosophila*. Of course, additional signals may also refine enhancer activity as JAK/STAT does, although candidate screens in our lab have yet to identify such a signal. Irrespective of other pathways, identifying JAK/STAT as a novel DR enhancer input helps to further define the regeneration enhancer code, which may aid in their identification, and that of gene regulatory networks (GRNs) related to injury repair in *Drosophila* and beyond. A notable example of a mapped regeneration GRN is in the acoel worm, where Egr acts as a master epigenetic regulator to active numerous genes in response to amputation injury (44). Here again, JNK signaling is implicated in activating genes proximal to damage-responsive regions regulated by Egr (44). Since DRs enhancers in imaginal discs are also epigenetically regulated (8, 16), it is possible that a similar regulatory situation might exist in which pioneer factors are required to regulate the regeneration program via DR enhancers. The best characterized pioneer factors in *Drosophila* are encoded by *grainy head* (*grh*) and *zelda* (*zld*), the latter of which has recently been shown to be important for maintaining cell fate identity in regenerating wing discs (81). Exploring their roles may further clarify how DR enhancers regulate regeneration in *Drosophila*.

### Establishing a JNK threshold necessary for DR enhancer activation

JNK signaling plays a central role in regeneration, with different levels or duration of signaling producing distinct outcomes (43, 63); low or transient JNK promotes proliferation and survival, while higher or sustained JNK leads to cell death (62). These outcomes may depend on whether signaling is autocrine or paracrine (43), and on the use of distinct JNK receptors. While the receptor Grindlewald (Grnd) is activated by the ligand Egr to trigger high JNK and cell death, Wengen (Wgn) is activated by ROS to promote cell survival and proliferation via P38 and potentially lower JNK levels (35). Thus, both signaling modality and receptor use may shape JNK’s effects.

Our data show that specific JNK activity levels determine downstream target gene expression. One open question is how high JNK can selectively induce JAK/STAT, enabling expression of targets like *wg* and *Mmp1* in areas where JNK alone is insufficient. An explanation could lie in the involvement of other factors. For example, P38 is activated alongside JNK in regenerating cells, which together drive JAK/STAT activity via *upd* expression (22). Alternatively, DR enhancers may regulate components of the JAK/STAT pathway themselves; indeed we identified damage-responsive regions adjacent to two *upd* genes (16), suggesting an autoregulatory loop of JAK/STAT signaling, akin to that of *dome* receptor feedforward expression observed in embryogenesis (82).

Although only recently discovered, DR enhancers have already revealed how regeneration-associated genes are activated, patterned and resolved. As our work here shows, these enhancers can be injury-specific and translate a single damage input into the spatially dynamic patterns of gene expression required for proper regeneration, shutting off once regeneration is complete. This makes them powerful tools to manipulate repair processes, an idea that has already led to methods driving expression of endogenous and exogenous factors to enhance regeneration in various organisms (8, 9), including adult mammals (10). Continuing the study of these enhancers is therefore essential to advance our understanding of regenerative biology.

## Materials and Methods

### Drosophila stocks

Flies were cultured in conventional dextrose fly media at 25°C with 12h light–dark cycles. Genotypes for each figure panel are listed in the <u>Supplementary file 1</u>. Fly lines used as ablation stocks are as follows: hs-FLP; hs-p65; salm-LexADBD, DVE>>GAL4 (DC^NA^), hs-p65/CyO; salm-LexADBD, DVE>>GAL4/ TM6B, Tb (DC^hepCA^)(16). Ablation stocks used for RNA-seq were previously outlined in (Harris et al. 2020). DR/DRMS reporters were previously described in (8, 16). The following stocks were obtained from Bloomington Drosophila Stock Center: AP-1-RFP (BL#59011), puc-lacZ (BL#11173), 10XSTAT-DGFP (BL#26200, BL#26199), 10XSTAT-GFP (BL#26197), UAS-Stat92E^RNAi^ (BL#33637), hh-GAL4 (BL#600186), Pnr-GAL4 (BL#3039), rn-GAL4 (BL#7405), Ptc-GAL4 (BL#44612), UAS-Hep^CA^ (BL#6406) UAS-Hep^WT^ (BL#3908), salm[GMR85E08]-GAL4 (BL#46804), mol-lacZ (BL#12173). UAS-mir(RHG) was a gift from the Iswar Hariharan at UC Berkeley (Siegrist et al. 2010), UAS-hop48a (UAS-hop) and UAS-hop^ORF^ were gifts from the Erika Bach at NYU. Stocks generated for this work: DR^WNT^ΔStat92E (II), DR^WNT^-RFP (II), DR^WNT^-GFP (III), DR^WNT^[NLS] (III). The

DR^Wnt^ΔSTAT92E transgenes were generated using In-Fusion PCR mutagenesis (Clontech, Mountain View, CA) to sequentially delete the consensus sequences (primers listed in <u>Supplementary file 1</u>). The DR^WNT^-RFP and DR^WNT^[NLS] transgenes were generated by replacing the *eGFP* coding sequence in the DR^WNT^-eGFP (Formerly, BRV-B-GFP (8) reporter construct with the *DSred* coding sequence (*Flybase id*:FBto0000019) from TRE-RFP (BDSC# 59011) (59), or Redstinger from pQUASp-RedStinger (nlsRFP) generated by Christopher Potter (Addgene plasmid # 46165). Cloning primers for constructs are listed in <u>Supplementary file 1</u>. The *hsp70* promoter used in all reporter constructs comprise the minimal promoter sequence with the heat shock elements removed (83) to prevent induction by the heat shock used in ablation experiments. Reporters were inserted into the *AttP40* landing site via PhiC31 recombination, DR^WNT^-eGFP (III) and DR^WNT^[NLS] (III). was inserted in the VK00027 (#BDSC 9744) cyto-location. Transgenic services were provided by BestGene (Chino Hills, CA)

### Ablation experiments

#### DUAL Control ablation

DUAL Control experiments were performed essentially as described in (16). Briefly, experimental crosses were cultured at 25°C and density controlled at 50 larvae per vial. Larvae were heat shocked on day 3.5 of development for early L3 (84 h after egg deposition [AED]) or day 4.5 for late L3 (108 h AED) by placing vials in a 37°C water bath for 45 min, followed by a return to 25°C. Larvae were allowed to recover for 18 h before being dissected, fixed and immunolabeled, unless otherwise indicated. All discs examined by immunofluorescence are early L3 discs ablated and imaged at the indicated hour AHS, with the exception of late L3 discs shown in Fig 1B. *UAS-y^RNAi^* were used as control lines for RNAi-based experiments.

#### GAL4/UAS ablation

GAL4/UAS-based ablation experiments were performed essentially as described in (16). Briefly, larvae bearing rn-*GAL4, GAL80^ts^, UAS-egr* were cultured at 18°C and density controlled at 50 larvae per vial. Larvae were upshifted on day 7 (168 h AED) or day 9 of development (216 h AED) for 40 h at 30°C and dissected into Trizol immediately and prepared for RNA-seq.

### Inducing JNK at different levels

Flies of genotype *UAS-hep^WT^, UAS-mir(RHG), salm[GMR85E08]-GAL4* were used for JNK^Low^ and JNK^Med^ conditions. Crosses were density controlled at 50 larvae per vial and kept at either 22°C (JNK^Low^) or 30°C (JNK^Med^) until mid L3 (120 h and 88 h AED) respectively before being dissected, fixed, and stained. Flies of genotype *UAS-hep^CA^, UAS-mir(RHG), GAL80^ts^, salm[GMR85E08]-GAL4* were used for the JNK^High^ condition. Crosses were density controlled at 50 larvae per vial and kept at 18°C until day 6 (144 h AED) when they were transferred to 30°C for 20 hours before being dissected, fixed, and stained.

### Immunohistochemistry

Larvae were dissected in 1 x PBS followed by a 20 min fix in 4 % paraformaldehyde in PBS (PFA). After 3 washes in 0.1 % PBST (1 x PBS + 0.1 % Triton-X), larvae were washed in 0.3% PBST and then blocked in 0.1 % PBST with 5 % normal goat serum (NGS) for 30 min. Primary staining was done overnight at 4°C, and secondary staining was done for 4 h at room temperature. The following primary antibodies were obtained from the Developmental Studies Hybridoma Bank: mouse anti-Nubbin (1:25), mouse anti-Wg (1:100), mouse anti-Mmp1 C-terminus (1:100), mouse anti-Mmp1 catalytic domain (1:100), mouse anti-LacZ (1:100), rat anti-Ci (1:3) and rat anti-DE-cadherin (1:100). Antibodies obtained from Cell Signaling Technologies: Rabbit anti-Dcp-1 (1:1000), mouse anti-PH3 (1:500). Secondary antibodies anti-rabbit 647, and anti-mouse 555 and anti-rat 555 were obtained from Invitrogen and used at a 1:500 dilution. DAPI (1:1000) was used as a counterstain. Images were obtained on a Zeiss AxioImager M2 with ApoTome. For each experiment at least 15 discs were analyzed prior to choosing a representative image, N used for quantification is indicated in figure legends. Images were processed using Affinity Photo and Affinity Designer. All images are set to the scale bar being 50 μm in length unless otherwise stated in the figure legend.

### DHE staining

Dihydroethidium (DHE) labeling was performed by incubating freshly dissected wing imaginal discs in Schneider’s Media with 1 µl of 10 mM DHE reconstituted in 1 ml DMSO (for a working concentration of 10 µm DHE) for 10 min, followed by three 5 min washes in PBS, fixed in 4% PFA for 8 mins, a further 1 min wash in PBS, followed by mounting and immediate imaging.

### Regeneration scoring and wing measurements

Wings of adult flies from heat shocked larvae were scored and measured after genotype blinding by another researcher. Scoring was performed on anesthetized adults by binning into a regeneration scoring category (16, 84). Wing measurements were performed by removing wings, mounting in Permount solution (Fisher Scientific) and imaged using a Zeiss Discovery V8 microscope. Wing area was measured using the Fiji software. Male and female adults were measured separately to account for sex differences in wing size using a reproducible measuring protocol that excludes the variable hinge region of the wing (details of measuring protocol available on request). Statistics were performed using GraphPad Prism 10.0.

### Quantification, Fluorescence Measurement and Statistical Analysis

Adult wings, mean fluorescence intensity, and cell counts were measured using Fiji. GraphPad Prism 10.0 was used for statistical analysis and graphical representation. Graphs depict the mean of each treatment, while error bars represent the standard deviation. The mean fluorescence intensity (MFI) was quantified in Fiji using arbitrary units (A.U). In each experiment, MFI of the wing pouch was normalized to the MFI of the non-fluorescent notum. For measurements of AP-1 intensity (Fig 1G), the profile tool on the Zen Imaging platform was used to generate data points across injured discs, which was standardized across disc widths by converting to percent distance, and analyzed in Prism. For measurement of STAT in the AP-1 domain (Fig 3G) the area of the AP-1 expression domain was selected and the MFI of 10XSTAT-DGFP in this domain was measured and normalized. For measurement of PH3 cell number (Fig 3H-I), cells were counted using the ImageJ cell counter plug-in. The wound center and periphery areas were selected by outlining the areas of high and low JNK expression using the AP-1 reporter, and the cells in each area counted. For all quantifications the mean and standard deviation for each normalized treatment was calculated and used for statistical analysis. The sample size and p values for all statistical analyses are indicated in the figure legends. Statistical significance was evaluated in Prism 10.0 using statistical test stated. Schematics were made using Biorender.com.

### qRT-PCR analysis

Total RNA was extracted from 15 wing discs per sample using the TRIzol reagent (Thermo Fisher Scientific), and cDNA libraries were prepared from total RNA using the AzuraQuant cDNA synthesis kit (Azuragenomics) according to manufacturers instructions. qRT-PCR was performed using AzuraQuant Green 1-step qPCR Mix LoRox (Azuragenomics) on a ThermoFisher Quant Studio3. Data were analyzed using the ΔΔCt method and normalized to the housekeeping gene actin5C. Primer sequences used for PCR: *Act5C* Fwd GGCGCAGAGCAAGCGTGGTA, Rvse GGGTGCCACACGCAGCTCAT; *DsRed* Fwd CACTACCTGGTGGAGTTCAAG, Rvse GATGGTGTAGTCCTCGTTGTG.

### RNA-seq

Library prep and RNA-Sequencing (RNA-Seq) was performed by the QB3 Genomics Facility at UC Berkeley from total RNA of ∼100 discs per sample using polyT adapters and sequenced on the Illumina Miseq platform.

#### Data processing

The RNA-Sequencing (RNA-Seq) data was processed using a pipeline that included quality control, trimming, read alignment, and gene expression quantification. Adapters and low-quality reads were trimmed using Trim Galore (bioinformatics.babraham.ac.uk/projects/trim_galore) using the Phred quality score of 20. We visualized quality of the raw and trimmed reads using FastQC and MultiQC (85). Trimmed reads were aligned to the *Drosophila melanogaster* reference genome obtained from FlyBase (Flybase.org; version r6.57) using HISAT2 (86). The resulting BAM files were sorted, and coverage statistics were calculated using SAMtools (87). Finally, gene-level read counts were obtained using featureCounts based on the *Drosophila melanogaster* annotation (version r6.57), also obtained from FlyBase (88).

#### Differential expression

Differential gene expression analysis was performed using the limma package in R (89) with the voom transformation (90) applied to account for the mean-variance relationship in RNA-seq data. After transforming the read counts into LOG_2_-counts per million (LOG_2_-CPM), a linear model was fitted to the data to estimate the differential expression between early and late L3 stages in damaged and undamaged conditions. Empirical Bayes moderation was applied to the standard errors to improve accuracy, and p-values were adjusted for multiple testing using the Benjamini-Hochberg method. Genes with an adjusted p-value less than or equal to 0.05 and an absolute LOG_2_ fold change (LOG_2_FC) greater than or equal to 2 were considered differentially expressed.

#### Gene enrichment analysis

A gene ontology (GO) enrichment analysis was performed using the clusterProfiler package in R (91) to identify the biological functions associated with the differentially expressed genes. The differentially expressed genes identified by the limma-voom analysis were used as input for GO enrichment analysis. Separate analyses were conducted for up-regulated and down-regulated genes between damaged and undamaged conditions in the early and late L3 stages. GO terms were considered significantly enriched if the adjusted p-value was less than or equal to 0.05.

### Transcription factor binding site analysis

The coordinates for regions classed as DR peaks (LOG_2_FC >0.5 and p adj <0.1) in either early or late L3 discs identified in Harris et al. 2020 were converted from dm3 to the current dm6 version r6.59 using the coordinates converter tool on Flybase (Flybase.org/convert/coordinates), and DNA sequences for each peak were extracted using the Extract Genomic DNA tool on Galaxy (usegalaxy.org) in FASTA format. The same process was used to obtain DNA sequences of static peaks, those that do not change between damaged and undamaged conditions (LOG_2_FC <0.5), which were used as the background sequences for TF enrichment analysis. The TF enrichment was performed using the AME tool on Meme Suite (meme-suite.org, (92) with default options, assaying for motifs in the Fly Factor survey database. Identification of Stat92E and AP-1 binding sites in DR regions was performed using the FIMO tool on Meme Suite with default settings and a match p-value of p<0.001 (meme-suite.org, (93)).

## Data and reagent availability

Stocks are available upon request and details of stocks and reagents used in this study are available in the materials and methods. The authors affirm that all data necessary for confirming the conclusions of the article are present within the article, figures, and tables. Raw data files will be available on SRA upon publication at Bioproject. The code for RNA-seq analysis is available here: https://github.com/mariahlee/Regeneration-in-Drosophila-melanogaster

## Acknowledgements

The authors would like to thank Dr. Iswar Hariharan and Dr Erika Bach for their generous gift of stocks and reagents. We thank the current members of the Harris lab for useful input and feedback. We thank the Bloomington Drosophila Stock Center and Developmental Studies Hybridoma Bank for stocks and reagents.

