## Supplementary figures and images for "A threshold level of JNK activates damage-responsive enhancers via JAK/STAT to promote tissue regeneration"

### Supplemental Figure 1

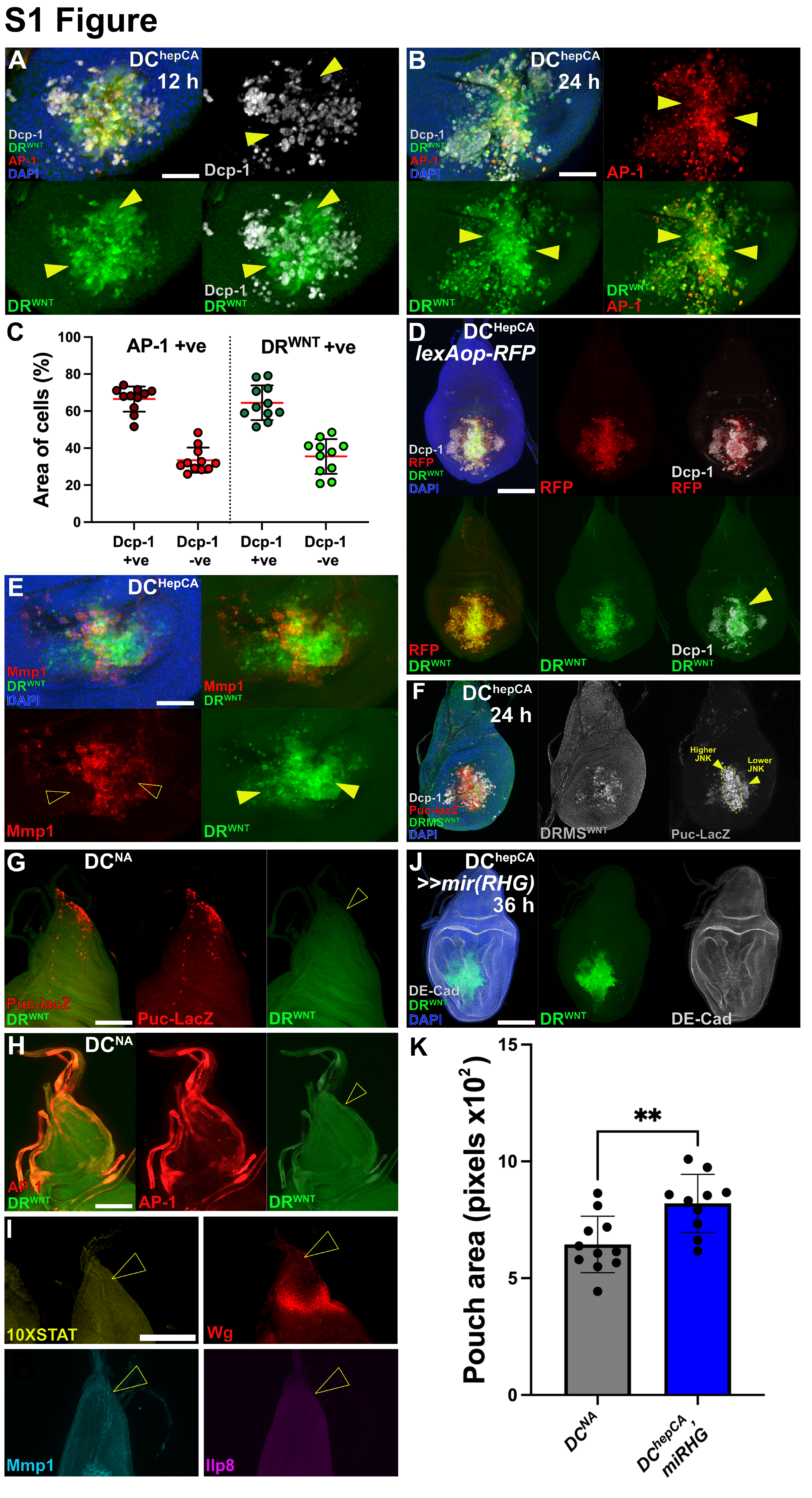

### Supplemental Figure 2

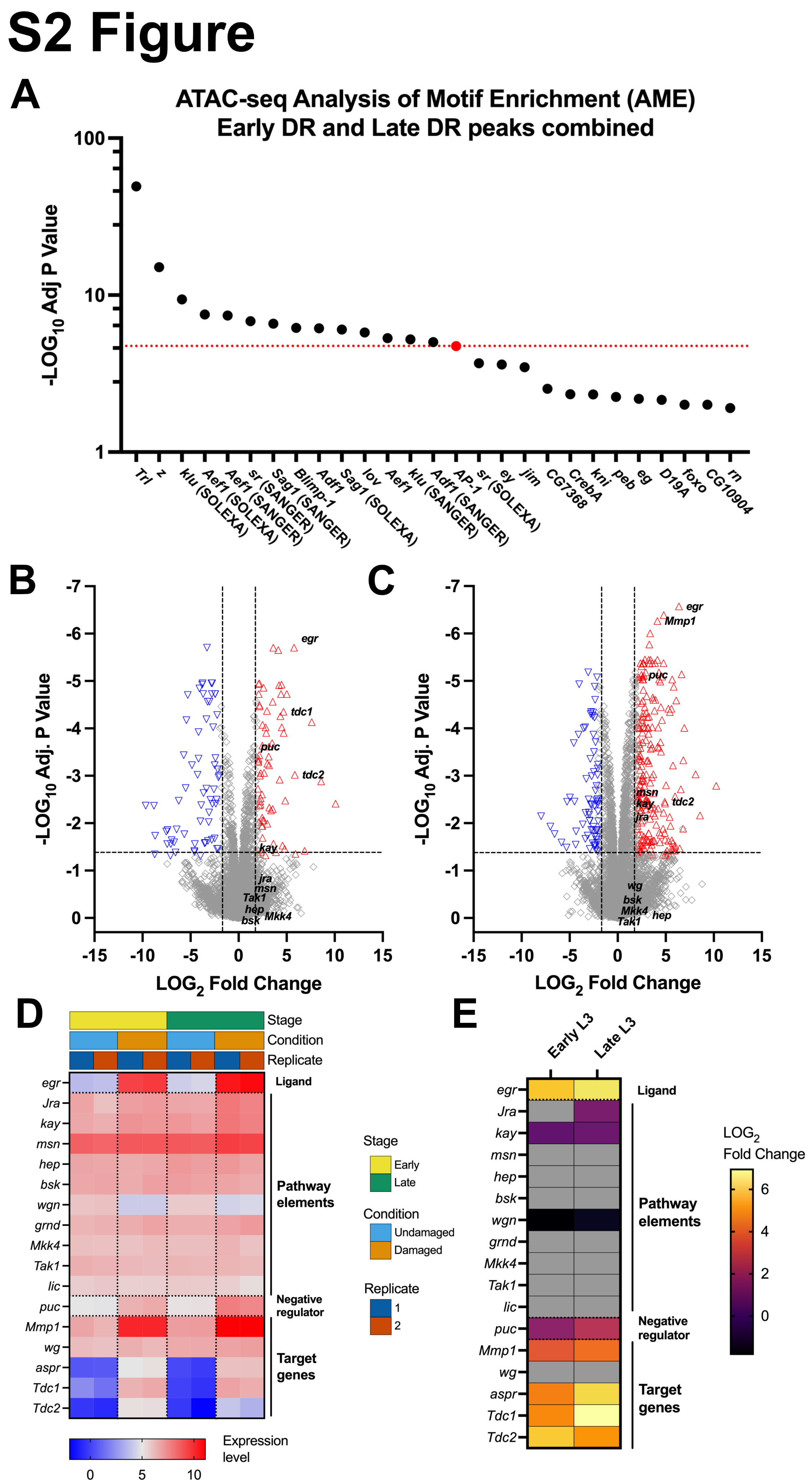

### Supplemental Figure 3

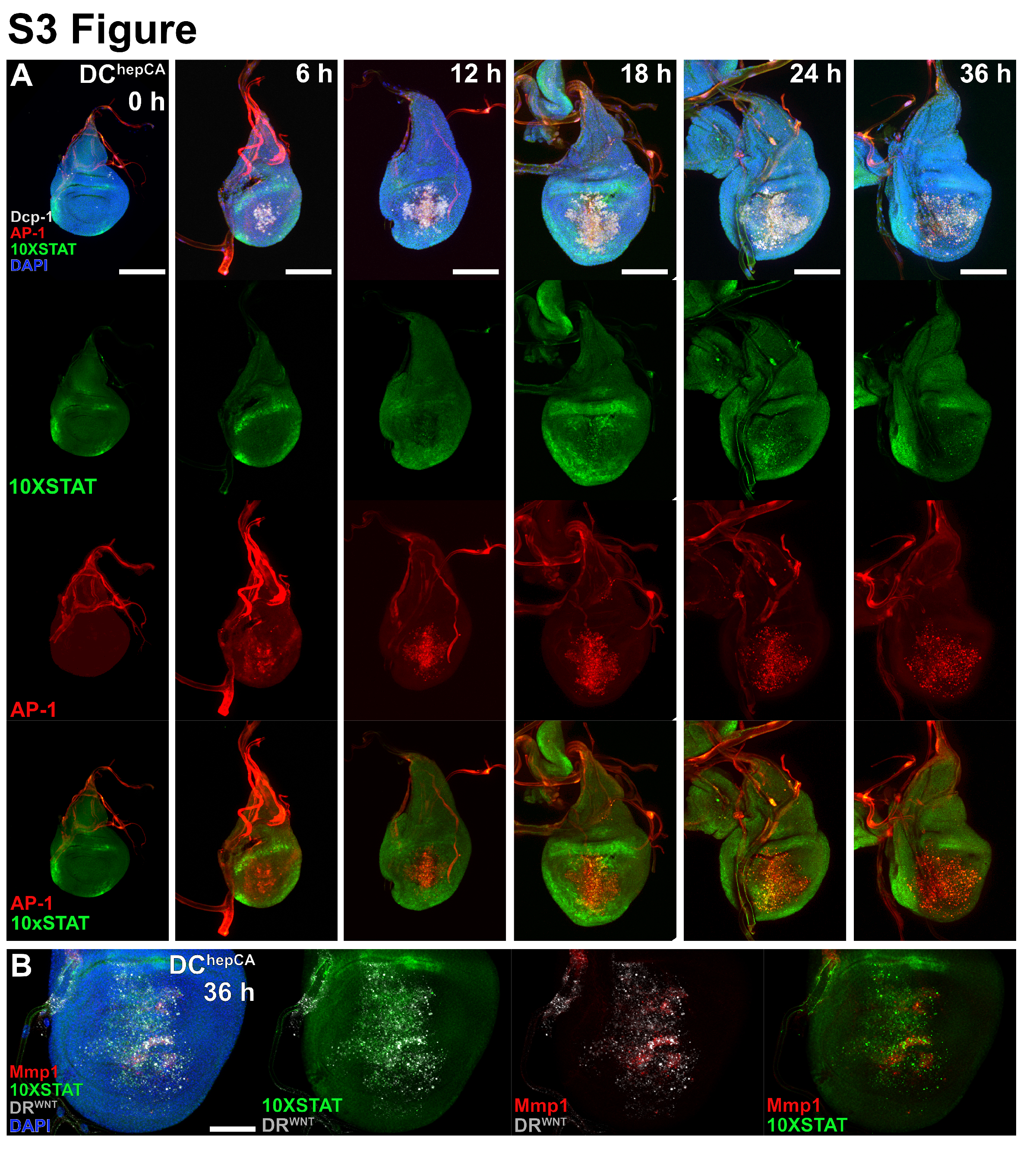

### Supplemental Figure 4

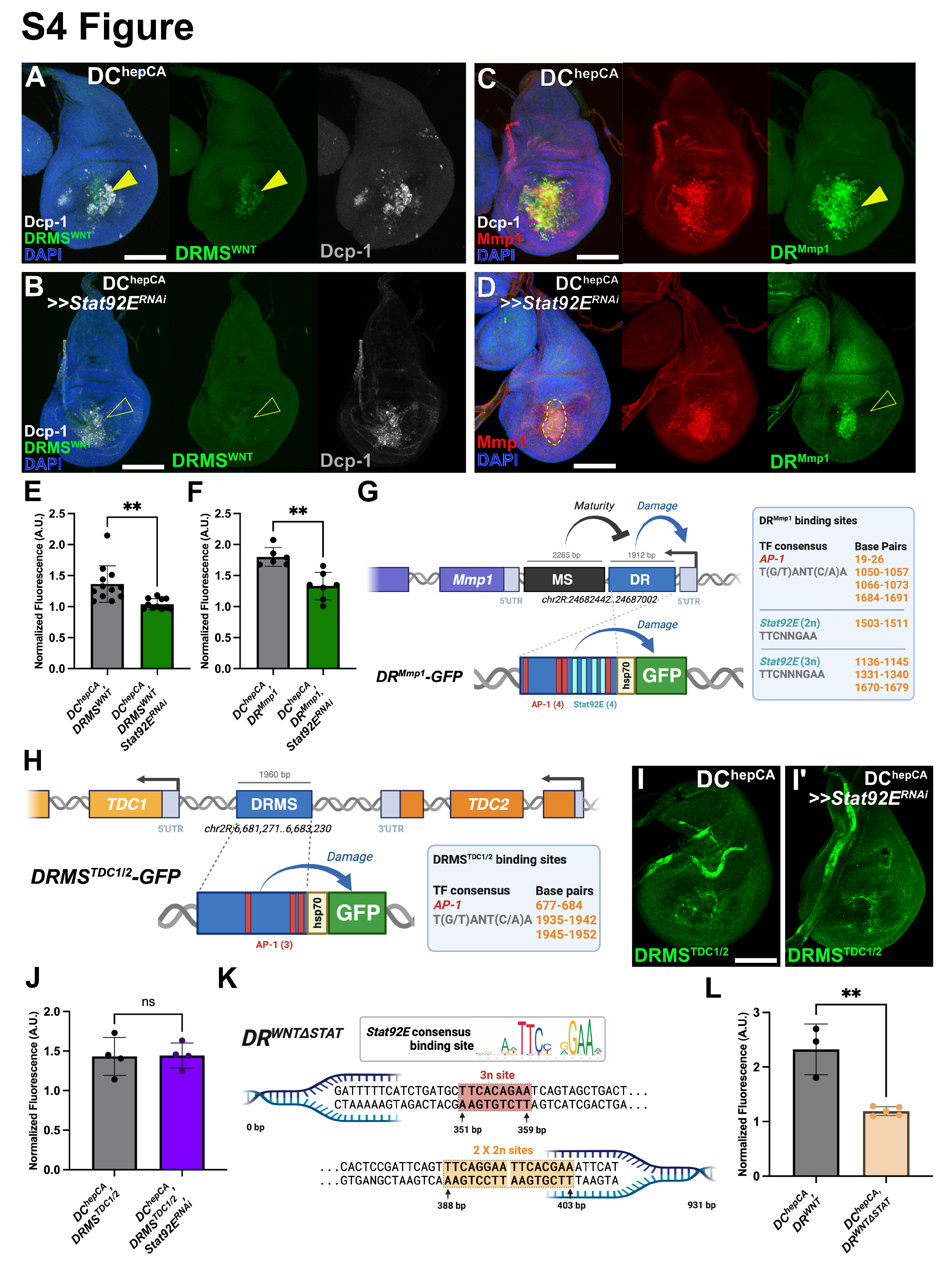

### Supplemental Figure 5

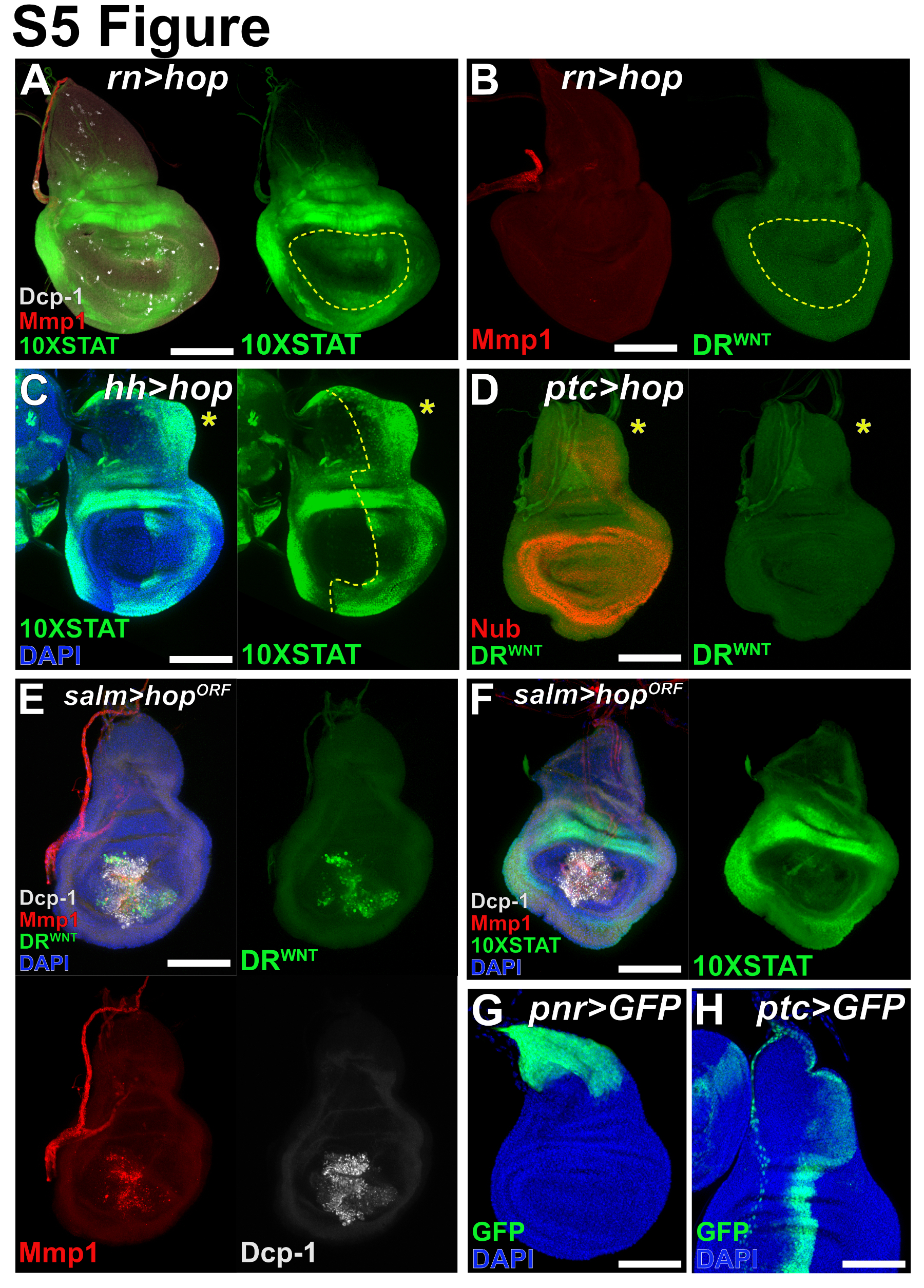

### Supplemental Figure 6

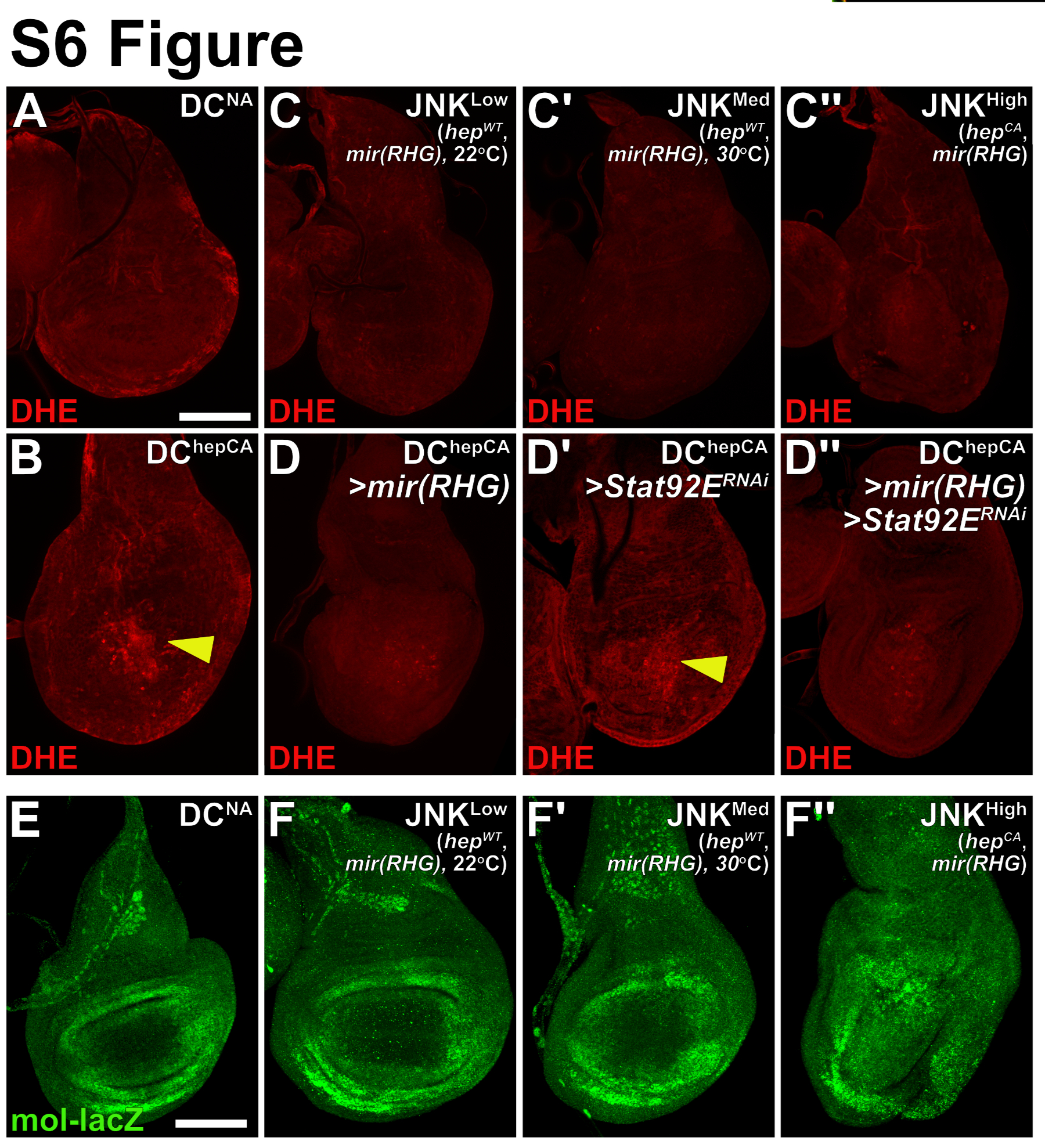
