## Supplemental Figure Legends for "A threshold level of JNK activates damage-responsive enhancers via JAK/STAT to promote tissue regeneration"

Supporting Information Captions

S1 Fig: Ablation activates DR^WNT^ and causes overgrowth in the absence of cell death.

**(A)** Disc bearing the non-NLS DR^WNT^ reporter (green) and AP-1 reporter (red) ablated by DC^hepCA^ at 12 h AHS, showing cells with DR^WNT^ activity (green) that lack Dcp-1 (white), indicated by yellow arrowheads. **(B)** Disc as in (A) at 24 h AHS, showing the DR^WNT^ reporter (green) does not entirely mirror the strength of the AP-1 reporter (red) across the disc, with differences in reporter expression indicated by yellow arrowheads. (**C**) Quantification of the number AP-1 positive cells or DR^WNT^ reporter positive cells that have Dcp-1 staining, shown as a percent of cell area with Dcp-1 staining. Error bars are SD. (**D**) Wing disc bearing the DR^WNT^ reporter (green), ablated by DC^HepCA^ and expressing *lexAop-RFP* (red) to label the ablation domain, stained with Dcp-1 (white) to show cell death. DR^WNT^ overlaps RFP, and ablation is consistent across the pouch. Yellow arrowheads indicate regenerating cells with DR^WNT^ activity and no Dcp-1. **(E)** Wing imaginal disc bearing the DR^WNT^ reporter (green), ablated by DC^HepCA^ and stained for Mmp1 (red). Yellow arrowheads and open arrowheads indicate cells with DR^WNT^ reporter activity that do not have Mmp1. **(F)** Disc bearing a reporter for the DRMS^WNT^ enhancer (green) and a reporter for JNK target *puc* (*puc-lacZ*, red), ablated by DC^hepCA^ and imaged at 24 h AHS. The *puc* reporter shows the separation of JNK activity into high and low levels (yellow arrowheads and dashed outline), and the activity of DRMS^WNT^ activity occurring in both regions, as for the DRWNT enhancer reporter. Dcp-1 staining indicates cell death (white). **(G-H)** Notum of undamaged discs bearing the DR^WNT^ reporter (green), with developmental JNK expression indicated by the JNK target *puckered* (*puc-lacZ*, red in G) or the AP-1 reporter (*AP-1-RFP*, red in I). The DR^WNT^ reporter is not activated by JNK signaling in the notum. **(I)** Notum area of the wing discs showing developmental expression of the 10XSTAT reporter, Mmp1, Wg and a Ilp8 reporter. Yellow arrowheads indicate expression, and an open arrowhead highlights the lack of expression. **(J)** JNK activity caused by DC^hepCA^ in the absence of cell death through mir(RHG) expression induces overgrowth, indicated by the DE-Cad (white). DRWNT reporter activity in green.

(**K**) Quantification of pouch size area (pixels x10^2^) using the tissue folds in the hinge as a marker for the pouch in non-ablated (DC^NA^) and ablated discs in which cell death is blocked (DC^hepCA^*,* mir(RHG)), showing mild but significant overgrowth. Error bars are SD. n = 11, (*DC^NA^*), n = 10 (*DC^hepCA^*, *mir(RHG)*), unpaired two-tailed t-test, **p = 0.004, Scale bars = 50 μm, except (A-B) and (G-H) = 25 μm.

S2 Fig: Whole genome analysis of early and late L3 discs identifies changes in factors associated with JNK signaling.

(**A**) Analysis of motif enrichment (AME) performed on DR regions identified in early L3 discs and late L3 discs by ATAC-seq. 243 DR regions were analyzed with 12,779 static peaks as control sequences. Enrichment is shown as -LOG10 adj p value. (**B-C**) Volcano plots of differentially expressed genes induced by damage identified by RNA-seq. LOG_2_ fold change (LOG_2_FC) vs. significance (-LOG_10_ adjusted p-value) of genes expressed in (C) early and (D) late L3 stages between damaged vs. undamaged conditions. JNK related genes are labelled. (**D**) Heatmap of the expression levels of JNK pathway genes across each stage (early vs late), damage condition (undamaged versus damaged) and replicate (first or second biological repeat), with red representing higher expression and blue representing lower expression. (**E**) Heatmap comparing damage-induced fold change (LOG_2_FC) of genes related to JNK signaling, including the ligands, pathway transducers, target genes and known negative regulators in early L3 versus late L3 discs. Gray cells indicate p-value is not significant (p>0.05).

S3 Fig: JNK and JAK/STAT signaling occur dynamically during regeneration.

(**A**) Full time course of discs ablated by DC^hepCA^ and imaged at 0, 6, 12, 18, 24, 36 h AHS during regeneration. Discs bear the 10XSTAT reporter of JAK/STAT activity (*10XSTAT-GFP*, green) and AP-1 reporter of JNK activity (*AP-1-RFP*, red). Cell death is labeled by Dcp-1 (white). **(B)** Disc bearing the 10XSTAT reporter (green) and DR^WNT^ reporter (cyan) ablated with DC^hepCA^, stained for Mmp1 and imaged at 36 AHS, showing the diminished expression of these factors in late regeneration. Scale bars in all panels = 50 μm, except (B) = 25 μm.

S4 Fig: Other DR/DRMS reporters are activated by damage, but not altered by JAK/STAT signaling.

(**A-B**) Discs bearing the DRMS^WNT^ reporter (*DRMS^WNT^-GFP*, green) ablated by DC^HepCA^, stained for Dcp-1 (white) and imaged at 18 h AHS, comparing wild type regeneration (A) to regeneration with knockdown of *Stat92E* (B). The lateral DRMS^WNT^ activity (yellow arrowhead in A) is reduced in ablated discs with *Stat92E* knockdown (open arrowhead in B). **(C-D)** Discs as in (A-B), but bearing the DR^Mmp1^ reporter (*DR^MMP1^-GFP*, green) and stained for Mmp1 (red) comparing wild type regeneration (C) to regeneration with knockdown of *Stat92E* (C). The lateral DR^Mmp1^ activity (yellow arrowhead in C) outside of the high JNK regions (yellow dashed outline in D) is absent from ablated discs with *Stat92E* knockdown (open arrowhead in D). **(E)** Quantification of DRMS^WNT^ reporter activity assayed by fluorescence intensity normalized to background in DC^hepCA^ ablated discs and DC^hepCA^ ablated discs with JAK/STAT blocked via *Stat92E* knockdown. Error bars are SD. n = 12 discs (DC^hepCA^), n = 11 discs (DC^hepCA^, *Stat92E^RNAi^*), unpaired two-tailed t-test, **p = 0.0021, A.U. = Arbitrary Units, **(F)** Quantification of DR^Mmp1^ reporter activity as in (E). Error bars are SD. n = 6 (DC^hepCA^), n = 7 (DC^hepCA^, *Stat92E^RNAi^*), unpaired two-tailed t-test, **p = 0.0011, A.U. = Arbitrary Units, **(G)** Schematic of the DRMS^Mmp1^ regulatory element (top), showing the separable damage-responsive (DR) region that activates *Mmp1* expression upon damage, and the maturity-silencing (MS) region that limits DR activity with developmental maturity. Flybase genomic coordinates (dm6) are shown below. The DR^Mmp1^ reporter (bottom), comprising the DR^Mmp1^ enhancer DNA, *hsp70* promoter and GFP coding sequence. Schematic shows AP-1 and Stat92E transcription factor binding sites in the *DR^Mmp1^* region, with details shown in table (right), including base pair range of binding sites within the genome coordinates. (**H**) Schematic of DRMS^TDC1/2^ labelled as in (G). This reporter has no sites matching the Stat92E consensus (**I**) Discs bearing the DRMS^TDC1/2^ reporter (*DRMS^TDC1/2^-GFP*, green) ablated by DC^hepCA^ with and without *Stat92E* knockdown, showing lack of activity in the periphery of the wound and no difference in damage-induced activity in the absence of JAK/STAT activity. (**J**) Quantification of DRMS^TDC1/2^ reporter activity assayed by fluorescence intensity normalized to background in DC^hepCA^ ablated discs and DC^hepCA^ ablated discs with JAK/STAT blocked via *Stat92E* knockdown. Error bars are SD. n = 4 (DC^hepCA^), n = 4 (DC^hepCA^, *Stat92E^RNAi^*), unpaired two-tailed t-test, ns = not significant, A.U. = Arbitrary Units, (**K**) Schematic showing the deletions of three potential Stat92E consensus binding sites to generate the DR^WNTΔSTAT^ reporter. Details are in materials and methods section. (**L**) Quantification of DR^WNTΔSTAT^ reporter activity compared to DR^WNT^ assayed by fluorescence intensity normalized to background in DC^hepCA^ ablated discs. Error bars are SD. n = 3 (DC^hepCA^, DR^WNT^), n = 5 (DC^hepCA^, DR^WNTΔSTAT^), unpaired two-tailed t-test, **p = 0.0014, A.U. = Arbitrary Units. Scale bars = 50 μm.

S5 Fig: JAK/STAT manipulation through *hop* expression.

**A)** Disc with ectopic JAK/STAT signaling in the distal pouch (dashed yellow outline) through overexpression of *hop* (*rn-GAL4*, *UAS-hop*), confirmed by activation of the 10XSTAT reporter (green). Dcp-1 (white) shows lack of cell death, Mmp1 (red) shows JNK activity **B**) Disc as in (A) showing the DR^WNT^ enhancer reporter (green) is not activated by ectopic JAK/STAT, **C**) Disc with ectopic JAK/STAT signaling in the posterior compartment (dashed yellow line shows A/P boundary) through overexpression of *hop* (*hh-GAL4*, *UAS-hop*), confirmed by activation of the 10XSTAT reporter (green). Asterisk highlights development of an ectopic pouch generated by JAK/STAT in the notum. (**D**) Disc with ectopic JAK/STAT signaling in the A/P boundary (dashed through overexpression of *hop* (*ptc-GAL4*, *UAS-hop*). Asterisk highlights development of an ectopic pouch generated by JAK/STAT in the notum, shown by nub staining (red). (**E**) Disc bearing the DR^WNT^ reporter (green) with high levels of JAK/STAT in the distal pouch through overexpression of the *hop^ORF^* transgene (*salm-GAL4, UAS-hop^ORF^*), which causes significant cell death, shown by Dcp-1 staining (white), and JNK activity shown by Mmp1 (red). (**F**) Disc as in (E) bearing the 10XSTAT reporter (green), showing *hop^ORF^* activates JAK/STAT signaling. (**G-H**) Discs with pannier-GAL4 (G) or patched-GAL4 (H) driving GFP (green) to demonstrate their expression domains (*pnr-GAL4, UAS-GFP*, or *ptc-GAL4, UAS-GFP*). Scale bar in all panels = 50 μm, except (H) = 35 μm.

S6 Fig: Production of ROS and expression of the ROS agonist *mol* following ablation and at different JNK levels.

(**A-D’’**) Visualization of reactive oxygen species (ROS) shown by DHE in discs of the indicated genotype/condition. Yellow arrowheads indicate higher levels of ROS. Non-ablated discs (DC^NA^) have no detectable ROS (A), while ablation with DC^HepCA^ leads to ROS production (B). ROS is not detected at any level of JNK activity when cell death is blocked by mir(RHG), which prevents JNK spreading (C-C’’). Discs ablated as in (B) but with cell death blocked by mir(RHG) limits ROS production (D). Blocking JAK/STAT signaling by *Stat92E* knockdown does not change ROS production (D’), while additionally blocking cell death by mir(RHG) does prevent ROS (D’’). (**E-F**) Discs bearing the *mol* reporter (*mol-lacZ*, green) in wild type conditions (DC^NA^, E) or in different levels of JNK activity (F-F’’), showing no change in reporter activity. Scale bars = 50 μm.

S1 Table: List of differentially expressed genes in damaged versus undamaged discs from early and late L3 larvae.

Genes that are differentially expressed upon damage in early and late L3 discs identified by RNA-seq. Data include unique Flybase gene ID, gene names and expression attributes.

S2 Table: Intersections of genes in damaged versus undamaged discs from early and late L3 larvae.

Grouping of genes showing their intersection across injury condition (damaged or undamaged) and developmental stage (early or late L3). Data include unique Flybase gene ID, gene names and expression attributes.

Supplementary File 1: Bloomington stock center numbers, genotypes of experiments in main figures, and primers used for cloning.
